# Layer 5 anterior cingulate cortical neurons engage dorsolateral periaqueductal gray excitatory neurons to facilitate the affective component of pain

**DOI:** 10.64898/2026.08.24.746788

**Authors:** Jasmine Kaslow, William M. McCallum, Amaury François, Gregory Corder, Eric J. Kremer, Kimberly Ritola, Nicole Mercer Lindsay, Grégory Scherrer

## Abstract

Pain is a conscious perceptual experience characterized by its aversive quality and consequent motivation to quench pain perception. The anterior cingulate cortex (ACC) critically contributes to the emotional dimension of pain. In both humans and rodents, ACC neural activity increases during acute and chronic pain, whereas ACC lesioning or excitability reduction decreases emotional reactivity during pain. However, the ACC is connected to many brain regions and is engaged during experiences beyond pain. Thus, it remains unclear through which circuit mechanisms the ACC shapes pain experience, and how specific those circuits are to nociception. Here, we show that excitatory input from the ACC to the dorsolateral periaqueductal gray (dlPAG) facilitates the affective-motivational dimension of pain. We first examined ACC→dlPAG connectivity using histology, optogenetics, and electrophysiology. We found that the axons of layer 5 ACC neurons terminate in the dlPAG and monosynaptically excite *Slc17a6+* (VGLUT2-expressing) dlPAG neurons. Second, we genetically targeted ACC→dlPAG neurons with viral vectors to express the inhibitory DREADD hM4Di and then exposed the animals to an array of pain tests. We found that, across acute and chronic pain states, inhibition of the ACC→dlPAG pathway reduced affective-motivational but not reflexive pain behaviors. Third, we used fiber photometry to record neural calcium activity in the ACC in behaving mice and found that ACC→dlPAG neurons are engaged during a broad array of aversive experiences, rather than exclusively during pain, and exhibit task-specific activity patterns. Collectively, these results uncover the direct contribution of ACC→dlPAG neural activity to pain unpleasantness and the necessity of this pathway for generating aversive behavioral responses in general, rather than specifically for encoding the unpleasant quality of noxious stimuli.

## Introduction

Pain is a sensory, emotional, and cognitive experience in which nociceptive stimuli trigger aversive reactions. Pain’s unpleasant quality is vital to motivate escape and learn to avoid harm, but can be debilitating in the absence of a true threat or when escape is impossible. Although the unpleasantness of pain holds significant potential for targeted pain relief strategies, the specific brain circuits that produce its affective component remain poorly understood.

Activity in the anterior cingulate cortex (ACC) has consistently been shown to correlate with pain affect. Human positron emission tomography (PET) and functional magnetic resonance imaging (fMRI) scans show increased signal in the ACC during unpleasant emotional states associated with diverse acute and chronic pain conditions (Apkarian et al., 2005; Rainville et al., 1997; Wager et al., 2013). In addition to recording noxious stimulus-evoked responses in the ACC of rodents (Kuo and Yen, 2005; Shyu et al., 2008; Wang et al., 2015, 2003; Wei and Zhuo, 2001), rabbits (Sikes and Vogt, 1992), and monkeys (Iwata et al., 2005), nociceptive single-units from intracranial microelectrode recordings have been identified in the human ACC (Hutchison et al., 1999). Patients with targeted surgical lesions of the cingulum bundle report their pain to be less distressing (Ballantine et al., 1967; Cohen et al., 2001; Foltz and White, 1962; Grahek, 2012). In rodents, ACC inhibition can reduce avoidance of cues associated with pain in conditioned place preference tests (Johansen et al., 2001; LaGraize et al., 2004), evoke preference for the ACC-inhibited chamber during chronic pain (Barthas et al., 2015; Li et al., 2010; Qu et al., 2011), and reduce spontaneous pain behavior (Gu et al., 2015). These studies suggest a necessary function of the ACC in processing the affective dimension of pain and, in particular, in linking an interoceptive state to motivated behavior. However, human imaging methods and whole-region lesions lack the cellular and anatomical specificity to draw precise conclusions about the circuit mechanisms by which ACC activity modulates pain experience (Journée et al., 2023).

Specifically, at least three critical questions remain unanswered (Kuner and Tan, 2021). First, the ACC projects broadly to multiple central nervous system (CNS) regions involved in generating emotions and pain perception, including the amygdala, striatum, periaqueductal gray (PAG), and spinal cord dorsal horn (Chen et al., 2018; Lee et al., 2022; Ma et al., 2022; Shi et al., 2022), and it is unclear which of these circuits are engaged by nociceptive ACC neurons to modulate pain experience. Second, there remains considerable debate regarding the nature of the ACC’s influence on the different dimensions of pain experience. A number of studies suggest that the ACC contributes exclusively to the affective component of pain (Barthas et al., 2015; Calejesan et al., 2000; Gu et al., 2015), while others report a modulatory effect on the sensory-motor features of pain experience, including nocifensive withdrawal reflexes (Acuña et al., 2023; Esmaeilou et al., 2022; LaGraize et al., 2004; Lee et al., 2022; Treede et al., 1999). Third, the ACC exerts a wide range of functions beyond encoding affect during the pain experience (Carter et al., 1999; Vogt, 2005). A fundamental question regarding the emotional aspect of pain is whether there exist dedicated nociceptive neurons within limbic circuits or, conversely, whether the same populations of ACC neurons encode affect during both pain and other emotional experiences.

To clarify these issues, we examined the connectivity, neural activity, and function of molecularly and anatomically defined nociceptive ACC neurons in pain. Using activity-dependent genetic labeling in TRAP2 mice, we identified a population of layer 5 *CamkIIɑ*+ ACC pyramidal neurons that are active during pain, which, unexpectedly, predominantly project to the PAG. We show that these cells engage excitatory neurons in the dorsolateral PAG columns, contrasting with classic ventrolateral PAG pain pathways that influence nociception at the spinal level (Basbaum and Fields, 1978) and with nociceptive ventrolateral PAG neurons that receive inputs from the prefrontal cortex (Huang et al., 2019), both of which influence nocifensive withdrawal responses. By manipulating their activity during pain, we demonstrate that ACC→dlPAG neurons selectively modulate affective-motivational pain behaviors without altering noxious stimulus detection or reflexive nocifensive behaviors. Furthermore, by performing optical recordings of ACC→dlPAG neuron activity in freely moving mice experiencing a variety of negative-and positive-valence experiences, we provide evidence that while these neurons are strongly responsive to noxious stimuli, their response profiles are neither nociception-nor salience-specific, arguing against a dedicated pain function. Because inhibition of ACC→dlPAG neurons is sufficient to profoundly reduce aversion associated with chronic pain, this nociceptive pathway may represent an attractive target for relieving pain via pharmacological intervention or neurostimulation.

## Results

### A nociceptive pathway from the ACC to the dorsolateral PAG columns

To identify ACC nociceptive neurons and their projections, we used genetic trapping in TRAP2 knockin mice (Fos^tm2.1(icre/ERT2)Luo^/J), which enables identification of neurons active during pain (Corder et al., 2019; DeNardo et al., 2019; Guenthner et al., 2013) (**Fig. S1**). In these mice, the neural-activity-dependent promoter of the *Fos* gene drives expression of the inducible DNA recombinase Cre-ERT2, and subsequent administration of the ERT2 ligand 4-hydroxytamoxifen (4-OHT) induces Cre activity. To trace the outputs of putative ACC nociceptive neurons, we injected an adeno-associated virus (AAV) AAVDJ-hSyn-FLEx-GFP-synaptophysin-mRuby into the ACC of TRAP2 mice, and three weeks later, injected 4-OHT and noxiously stimulated the plantar surface of their hindpaw with pinpricks. This procedure labeled putative nociceptive neurons in the ACC with somatic GFP and presynaptic synaptophysin-mRuby. Examination of subcortical regions for synaptophysin-mRuby+ axon terminals indicated that ACC nociceptive neurons project densely to multiple structures with known function in pain affect, such as the basolateral amygdala (BLA), nucleus accumbens (NAc), and periaqueductal gray (PAG), as well as more sparsely to regions associated with modulating nocifensive reflexes like the rostral ventromedial medulla (RVM) (**Fig. S1**).

With respect to pain, the PAG is best known for its ventrolateral columns (vlPAG), which contain descending pain-modulatory neurons that regulate nociception at the spinal level via the RVM (Bagley and Ingram, 2020; Livrizzi et al., 2026). However, when visualizing the distribution of synaptophysin-mRuby+ terminals within the different columns of the PAG, we found that putative nociceptive neurons in the ACC predominantly project to the dorsolateral columns (dlPAG) (**Fig. S1**). To investigate the properties of these ACC→dlPAG neurons, we next used a dual viral targeting approach in wildtype mice, injecting a Cre-expressing virus with retrograde transport properties (AAV2-retro-Cre or CAV-Cre (Del Rio et al., 2019; François et al., 2017; Tervo et al., 2016)) into the dlPAG, and an AAV-Promoter-FLEx-transgene into the ACC. When we injected an AAVDJ-hSyn-FLEx-GFP-synaptophysin-mRuby into the ACC (**Figs. 1A,B, S2**), we observed that GFP/mRuby fluorescence predominated in layer 5 pyramidal (L5p) ACC neurons, identified by their location and morphology, with dendrites spanning all superficial layers (**Fig. 1B, left**) and branching apical tufts, and confirm their fibers are present the dlPAG (**Fig. 1B, right**), consistent with our anatomical findings in TRAP2 mice. Notably, we observed an absence of GFP and mRuby signal in the BLA (**Fig. 1C, left**), which is essential for pain emotional responses; nor in the spinal cord dorsal horn (**Fig. 1C, right**), where inputs from the ACC facilitate nociception (Chen et al., 2018, 2014). Instead, mRuby+ axon terminals were present in the medial thalamus, zona incerta, and pontine nucleus (**Fig. S2**), suggesting these axons represent collaterals of L5p ACC→dlPAG neurons, rather than outputs of separate populations of ACC neurons.

**Figure 1.**
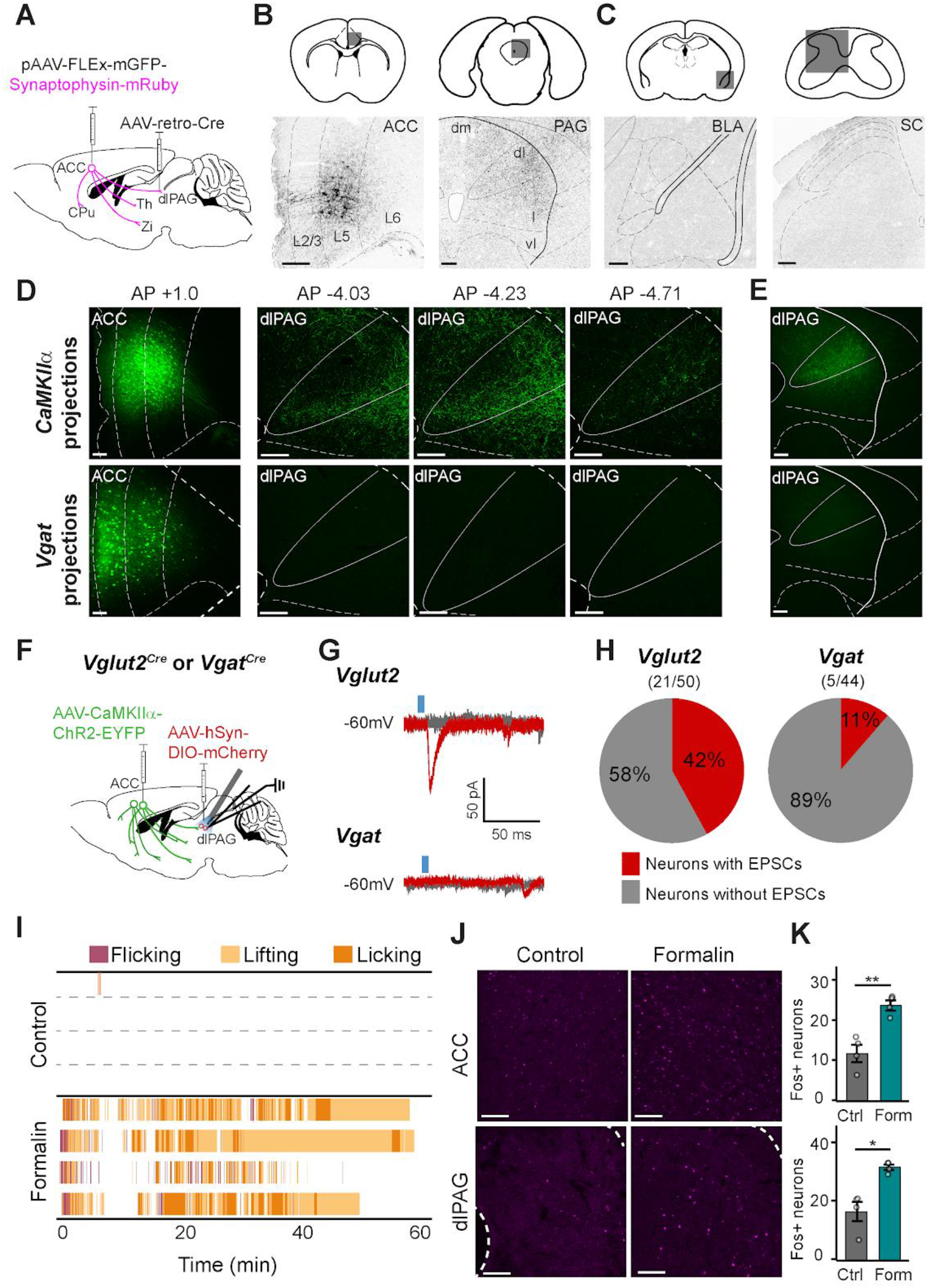
ACC projections to dorsolateral PAG. **A)** Virus injection schematic for tracing ACC→dPAG neuron axon collaterals. **B)** GFP expression in L5 ACC→dlPAG cell bodies (*Left*) and axons in dlPAG (*Right*). **C)** Absence of off-target axons from L5 ACC→dlPAG neurons in the Basolateral Amygdala (BLA, *Left*) and lumbar spinal cord dorsal horn (SC, *Right*). **D)** ACC *CaMKIIα* excitatory cell bodies and axons projecting to the dlPAG at three different anterior-posterior coordinates (*Top*) and ACC *Vgat* inhibitory cell bodies and lack of axons projecting to the dlPAG (*Bottom*). **E)** ACC *CaMKIIα* excitatory neuron axons projecting to the dorsal, but not ventral PAG (*Top*). Absence of ACC EYFP-labeled inhibitory neuron axons projecting to the whole PAG (*Bottom*). **F)** Virus injection and electrophysiology recording schematic for testing monosynaptic ACC inputs to dlPAG neurons. **G)** Representative excitatory postsynaptic current from *Vglut2* (*Top*) or *Vgat* (*Bottom*) dlPAG neurons in response to ACC ChR2 excitation. **H)** Proportion of neurons within each genetically defined dPAG population receiving monosynaptic inputs from the ACC. **I)** Raster plot of behavioral epochs after either no injection or an injection of 10 μL of 5% formalin in the left hindpaw. Each row represents one mouse. **J)** FOS in the ACC or dlPAG 90 minutes after hindpaw formalin injection. **K)** Mean FOS cell counts per animal for the ACC (*Top*) or dlPAG (*Bottom*). *p < 0.05, **p < 0.01. All scale bars = 100 µm. All error bars indicate SEM.

Next, to confirm that L5p ACC→dlPAG neurons are excitatory, we selectively labeled either ACC excitatory pyramidal neurons by injecting an AAV5-CamkIIɑ-FLEx-YFP in the ACC of wildtype mice, or ACC inhibitory neurons by injecting an AAV5-hSyn-FLEx-GFP in the ACC of *Slc32a1*^Cre^ mice (*i.e.*, with Cre expression in VGAT+ neurons), and traced their axons in the PAG (**Fig. 1D,E**). YFP+ axons from ACC excitatory pyramidal neurons showed a similar projection pattern to the ACC→dlPAG neurons (**Figs. 1A-C, S2**), with axon terminals in the dlPAG columns (**Fig. 1D,E, upper**). In contrast, we observed no long-distance projections from ACC inhibitory neurons to the PAG (**Fig. 1D,E, lower**).

We next tested functional connectivity between the ACC and dlPAG. We used whole-cell patch-clamp electrophysiology and optogenetics in brain slices to test monosynaptic connectivity between ACC and dlPAG neurons and to identify properties of PAG neurons that receive ACC inputs (**Fig. 1F**). We injected an AAV5-CamkIIɑ-ChR2-YFP in ACC and an rAAVDJ-EF1ɑ-DIO-mCherry in dlPAG in *Slc32a1*^Cre^ or *Slc17a6*^Cre^ mice (*i.e.*, with Cre expression in VGAT+ or VGLUT2+ neurons, respectively). After shining brief pulses of blue light onto the slice while recording from the mCherry+ PAG neurons, we observed light-evoked excitatory postsynaptic currents (EPSCs) with constant latency and no failures in neurons located in dlPAG, indicating their monosynaptic connection from ChR2-YFP+ L5p ACC neurons (**Fig. 1G**). Furthermore, the great majority of these monosynaptic inputs were recorded in slices from *Slc17a6*^Cre^ mice (21/50 neurons, 2-proportion Z-test, p = 0.00094) and were seldom observed in slices from *Slc32a1*^Cre^ mice (5/44 neurons) (**Figs. 1H, S3A**), revealing a putative nociceptive excitatory circuit that links the ACC and the dlPAG.

To identify whether ACC and dlPAG neurons show similar responses to pain experience, we injected formalin into the hindpaw, a pronociceptive insult that induces pain for ∼1 hour (**Fig. 1I**) (Tjølsen et al., 1992). As expected, we observed both reflexive and affective motivational pain behaviors in formalin-injected but not control mice (**Fig. 1l**). Notably, we identified concurrent increases in Fos+ neurons in the ACC and dlPAG compared to control mice, implicating both regions in nociceptive processing (**Figs. 1J,K, S3B,C**) (ACC: unpaired t-test, p = 0.005479; dlPAG: unpaired t-test, p = 0.01538).

### Inhibiting L5p ACC→dlPAG neurons decreases acute affective-motivational pain behaviors without altering noxious stimulus detection or nocifensive reflexes

The precise contribution of ACC neurons to pain experience remains a topic of debate. Specifically, rodent pain studies in which the ACC was lesioned, or its activity was altered, have reported changes either exclusively in affective-motivational behaviors (Barthas et al., 2015; Calejesan et al., 2000; Gu et al., 2015; Oswell et al., 2026), exclusively in nocifensive withdrawal reflexes, or in both (Acuña et al., 2023; Esmaeilou et al., 2022; LaGraize et al., 2004; Treede et al., 1999). Given the considerable diversity of cortical neurons, including L5p neuron types (Baker et al., 2018; BRAIN Initiative Cell Census Network (BICCN), 2021; Tasic et al., 2018), these divergent conclusions may stem from manipulations of distinct ACC neuronal populations.

To determine the function of L5p ACC→dlPAG neurons in pain, we inhibited these cells using chemogenetic DREADDs (Roth, 2016) and examined the effect of this pathway-specific inhibition on nocifensive reflexes and affective-motivational behaviors. To express the inhibitory DREADD hM4Di in L5p ACC→dlPAG neurons, we bilaterally injected a rAAV-DIO-hM4Di-mCherry into the ACC and a CAV-Cre into the dlPAG of wildtype mice (**Fig. 2A**). This strategy yielded successful expression of hM4Di-mCherry in L5p cells along the ACC’s rostral-caudal axis, with peak expression at approximately +0.9 mm anterior to bregma (**Fig. 2B,C**).

**Figure 2.**
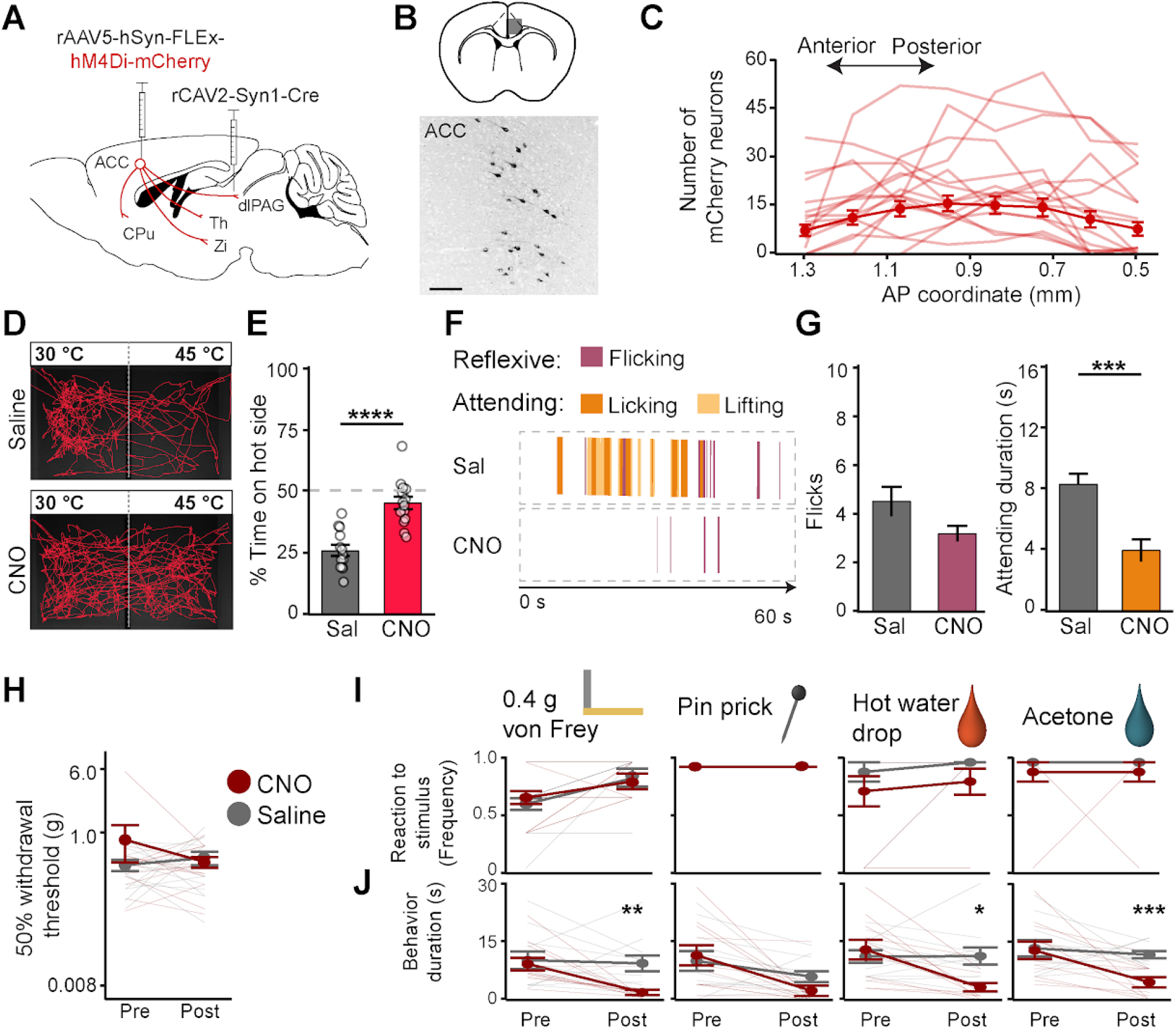
Chemogenetic inhibition of ACC→dlPAG neurons abolishes affective-motivational but not reflexive pain responses. **A)** Virus injection schematic for selectively inhibiting ACC→dlPAG projection neurons. All injections were bilateral. **B)** hM4Di-mCherry expression in L5 ACC neurons. **C)** Distribution of bilateral hM4Di-mCherry neuron counts across the rostral-caudal axis of the ACC. Thin lines represent individual animals, with the thick line indicating the population means. **D)** Representative movement trace of hM4Di-mCherry mice with saline vehicle or CNO during the thermal place avoidance test. **E)** Mean time spent on the hot side for mice injected with either saline vehicle or CNO. **F)** Representative raster plots of behavioral epochs during the 60-second hot plate test. **G)** Mean number of reflexive flicks for each test group on the hot plate (*Left*) and mean duration of attending behavior (*Right*). **H)** von Frey up-down mechanical threshold test pre and post CNO or saline vehicle. **I)** Frequency of reflexive hindpaw responses to each punctate stimulus. **J)** Duration of attending (prolonged lifting, licking, grabbing, biting) and escape (jumping, rapid movement) behavior in the 30-second window following stimulus application. *p < 0.05, **p < 0.01, ***p < 0.001. All scale bars = 100 µm. All error bars indicate SEM.

First, in a thermal place avoidance test in which adjacent plates were set to 45°C and 30°C, we found that mice injected with Clozapine-N-Oxide (CNO, to induce hM4Di-mediated inhibition in L5p ACC→dlPAG neurons) showed reduced avoidance of the noxiously hot side compared to control mice (**Figs. 2D,E, S4A-C**, unpaired t-test, p < 0.0001, n = 13 saline, 14 CNO), without observable changes in locomotion (**Fig. S4D,E**). We next employed a modified hot plate test (Corder et al., 2017), in which we recorded all nocifensive behaviors displayed over 60 seconds and categorized them as either reflexive or affective-motivational pain behaviors (Corder et al., 2019, 2017; Manglik et al., 2016). While we observed no change in the number of reflexive paw flicks (unpaired t-test, p = 0.06446), L5p ACC→dlPAG inhibition significantly decreased affective-motivational behaviors such as paw licking and guarding (**Figs. 2F,G, S5a**, unpaired t-test, p < 0.001, n = 15 saline, 16 CNO). The latency to first response for either behavioral category remained unchanged in both groups (**Fig. S5B,C**), suggesting intact sensitivity and reflexive motor responses.

To further examine mouse pain sensitivity and affect, we compared the reflexive withdrawal threshold with behavioral responses to somatosensory stimuli of different modalities applied to the hindpaw: von Frey filament, pinprick, hot water drop (noxious heat), or acetone (innocuous cool). For the latter test, similar to the hot plate assay in **Fig. 2F,G**, we scored both reflexive reactions and affective-motivational responses by identifying both the frequency of the reflex response and the duration of affective-motivational behavior following the stimulus (*e.g.,* such as nocifensive behaviors directed to the stimulus site, rapid general movement, and/or escape seeking, in the 30-second window after stimulus application).

ACC→dlPAG inhibition did not alter the mechanical withdrawal threshold compared to saline controls (**Fig. 2H**, 2-way repeated measures ANOVA; p = 0.5671 with respect to treatment, p = 0.2405 with respect to time point × treatment interaction). Between CNO-and saline-treated mice, we observed no difference in frequency of reflexive responses for any of the somatosensory stimuli we applied (**Fig. 2I**, 2-way repeated measures ANOVA for each; 0.4 g hair: p = 0.8704 with respect to treatment, p = 0.4947 with respect to time point × treatment interaction; pinprick: equivalent responses for all mice across all groups; hot water: p = 0.1828 with respect to treatment, p = 1.0000 with respect to time point × treatment interaction; acetone: p = 0.1727 with respect to treatment, p = 1.000 with respect time point × treatment interaction). In sharp contrast, we found that CNO-injected mice showed a decrease in the duration of affective-motivational behaviors exhibited after stimulus onset (**Fig. 2J**, 2-way repeated measures ANOVA for each; 0.4 g hair: p = 0.0419 with respect to treatment, p = 0.0513 with respect to time point × treatment interaction, post-drug time point, p = 0.0096 with respect to treatment group; pinprick: p = 0.6287 with respect to treatment, p = 0.1919 with respect to time point × treatment interaction; hot water: p = 0.1886 with respect to treatment, p = 0.0040 with respect to time point × treatment interaction, post-drug time point, p = 0.0110 with respect to treatment group; acetone: p = 0.0402 with respect to treatment, p = 0.0746 with respect to time point × treatment interaction, post-drug time point, p = 0.00061 with respect to treatment group; n = 14 saline, 14 CNO, Benjamini-Hochberg multiple comparison’s correction). We confirmed that this change in nocifensive behavior was not due to decreased general movement, as CNO-and saline-injected mice demonstrated similar distances traveled and velocities in the open field test (**Fig. S6**). Finally, to ensure that behavioral changes were not due to an off-target effect of CNO, we tested the effects of CNO in wildtype mice with no hM4Di-mCherry expression and saw no effect in the thermal place avoidance test or in their distance traveled or velocity in the open field test (**Fig. S7**).

Altogether, these results show that L5p ACC→dlPAG neurons are required for nocifensive affective-motivational behaviors, but do not influence reflexive withdrawal from noxious stimuli.

### L5p ACC→dlPAG neurons mediate the full expression of chronic pain-associated negative affect

Chronic pain states generally include pathological perceptual changes. Patients with peripheral neuropathic pain often suffer from hypersensitivity to cold and mechanical stimuli (Baron, 2000). Thus, we next asked whether L5p ACC→dlPAG neurons mediate affective-motivational behaviors associated with neuropathic hypersensitivity, using the spared nerve injury (SNI) model in mice expressing hM4Di-mCherry in these neurons (**Fig**. **3A**).

**Figure 3.**
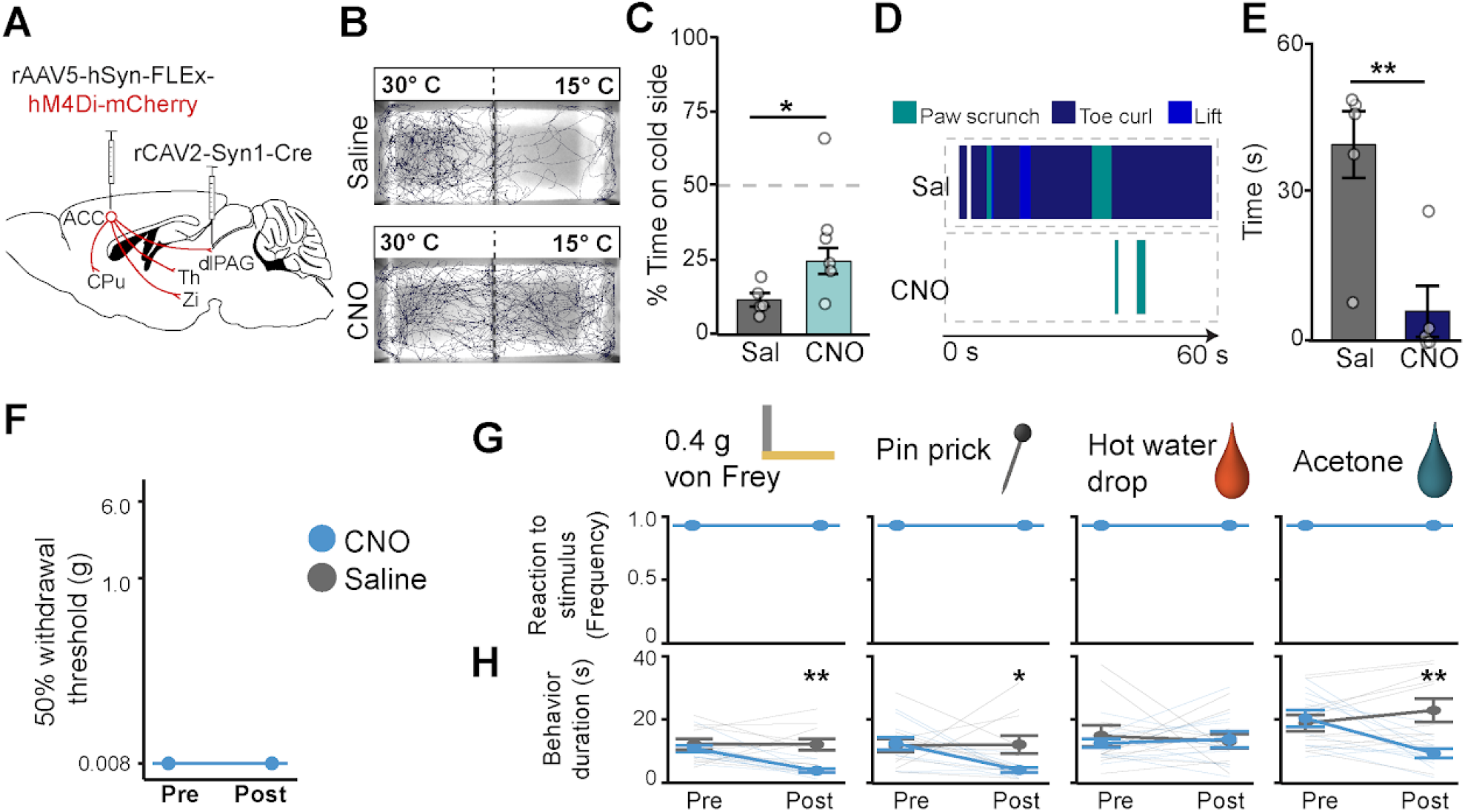
Chemogenetic inhibition of ACC→dlPAG neurons ameliorates affective-motivational nocifensive behaviors associated with chronic pain. **A)** Virus injection schematic for selective inhibition of ACC→dlPAG projection neurons. All injections were bilateral. **B)** Representative movement trace of hM4Di-mCherry mice during the cold place avoidance test after injection of saline vehicle or CNO. **C)** Mean time spent on the cold plate side for mice injected with either saline vehicle or CNO. **D)** Representative raster plots of behavioral epochs during the 60-second cold plate test. **E)** Mean duration of behavioral engagement for the cold plate test. **F)** von Frey up-down mechanical threshold test pre and post CNO or saline vehicle. **G)** Frequency of reflexive hindpaw responses to each punctate stimulus. **H)** Duration of attending (prolonged lifting, licking, grabbing, biting) and escape (jumping, rapid movement) behavior in the 60-second window after stimulus application. *p < 0.05, **p < 0.01. All scale bars = 100 µm. All error bars indicate SEM.

To assay affective-motivational behaviors associated with cold hypersensitivity, we first used the thermal place avoidance test, with one plate set at 15 °C and the other at 30 °C (**Fig. 3B**). In this assay, mice with SNI show strong avoidance of the colder plate (Corder et al., Science, 2019). Here, CNO-injected mice exhibited both reduced cold avoidance as well as increased distance traveled and velocity in the open field compared to saline-treated mice (**Figs. 3B,C, S8A-E**, unpaired t-test, p = 0.02298, n = 7 saline, 6 CNO), suggesting L5p ACC→dlPAG neurons influence neuropathic cold hypersensitivity.

We next modified the hot plate test to capture cold-induced pain affective-motivational behaviors in neuropathic animals. When placed on the cold plate for 60 seconds, SNI mice displayed a markedly different profile of discomfort-related behaviors compared with the hotplate test. Instead of predominantly large-limb or whole-body relief efforts, SNI mice displayed more subtle affective-motivational behaviors such as scrunching and tilting the injured paw away from the floor (**Figs. 3D, S9A**). These behaviors were largely absent in uninjured mice, suggesting that they result from neuropathic cold hypersensitivity (**Figs. 3D,E, S9B,C**). Notably, when we assayed the withdrawal threshold, mice injected with CNO showed unperturbed reflex responses (**Figs. 3F, S8F**; unpaired t-test, p > 0.05, n = 6 saline, 5 CNO), implying that inhibition of L5p ACC→dlPAG neurons impacts only pain affect.

To further examine the contribution of L5p ACC→dlPAG neurons to neuropathic hypersensitivity, we scored responses of SNI mice to cool (acetone) and mechanical (von Frey filament and pinprick) stimuli applied to the injured hindpaw. CNO-mediated inhibition of L5p ACC→dlPAG neurons did not alter the frequency of nocifensive responses to these stimuli (**Fig. 3G**). However, for von Frey, pinprick, and acetone, we found that CNO significantly decreased the duration of affective-motivational behaviors exhibited following stimulus application (**Fig. 3H**, 2-way repeated measures ANOVA for each; 0.4 g hair: p < 0.01 with respect to treatment, p < 0.01 with respect to time point × treatment interaction, post-drug time point, p < 0.01 with respect to treatment group; pinprick: p = 0.0882, with respect to treatment, p < 0.05 with respect to time point × treatment interaction, post-drug time point, p < 0.05 with respect to treatment group; Hot water: p = 0.7273, with respect to treatment, p = 0.5704 with respect to time point × treatment interaction, acetone: p = 0.1009, with respect to treatment, p < 0.001 with respect to time point × treatment interaction, post-drug time point, p < 0.01 with respect to treatment group; n = 14 saline, 14 CNO).

Our results thus far reveal that L5p ACC→dlPAG neurons represent an essential neural substrate for the full expression of pain affect during both physiological nociception and chronic pain.

### L5p ACC→dlPAG neurons broadly respond to noxious and aversive somatosensory stimuli

The ACC subserves a wide range of functions beyond encoding pain affect (Vogt, 2005). A fundamental question within limbic and cognitive circuits is whether cell populations are specifically dedicated to pain experience or, alternatively, whether the same cells that encode non-nociceptive experiences also mediate pain affect. One barrier to answering this question has been the limited data on the population dynamics of individual L5p ACC neuron types. Therefore, to assess how L5p ACC→dlPAG neurons contribute to nociception relative to the broader ACC pyramidal population—and to distinguish this from general valence or salience encoding—we next performed calcium imaging in freely moving mice.

To compare neural activity in L5p ACC→dlPAG with that of general excitatory ACC neurons, we expressed the genetically encoded calcium indicator GCaMP6s and recorded population transients using fiber photometry. First, we injected either AAV2-retro-GCaMP6s into the dlPAG or AAV-CamkIIɑ-GCaMP6s into the ACC and subsequently placed the recording fiber in the ACC (**Figs. 4A, S10**). To characterize the response properties of these two ACC neuron populations, we applied an assortment of stimuli (noxious heat, pinprick, acetone, von Frey filament, air puff, and approach-miss as a control) to the mouse hindpaw while recording (**Fig. 4B**).

**Figure 4.**
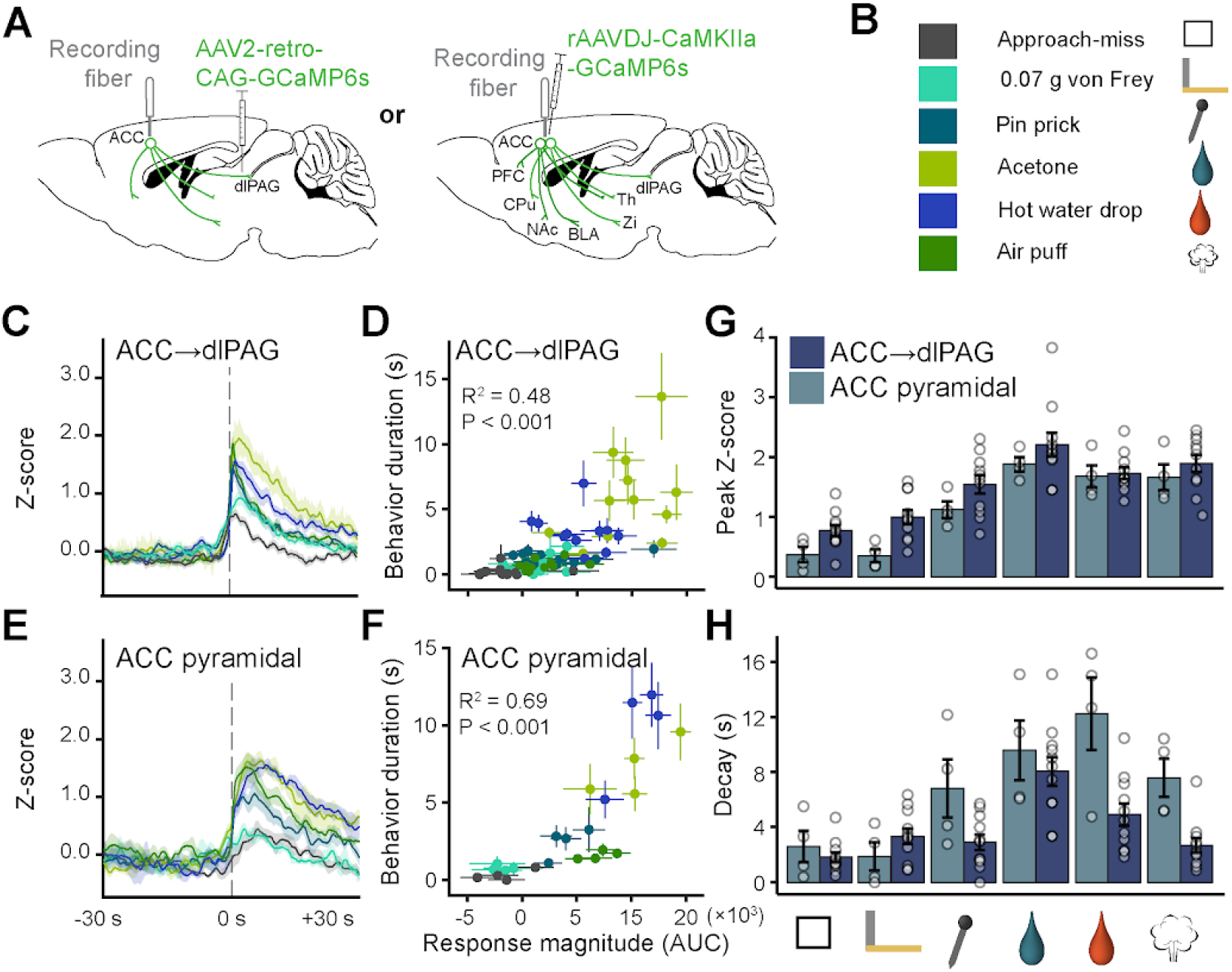
ACC→dlPAG neurons sharply encode aversive experience. **A)** Virus injection schematic to selectively record activity of either ACC→dlPAG projection neurons (*Left*) or all ACC excitatory pyramidal neurons (*Right*). Injections and implants were unilateral. **B)** Description of stimuli with color code and icon legend. **C)** ACC→dlPAG mean fluorescence trace ± SEM during stimulus application. **D)** Pearson correlation of ACC→dlPAG response magnitude and behavior duration in the 30-second post-stimulus window. One dot = one mouse. Error bars reflect SEM for individual trials on the respective axis. **E)** ACC broad excitatory population mean fluorescence trace ± SEM during stimulus application. **F)** Pearson correlation of ACC broad excitatory response magnitude and behavior duration in the 30-second post-stimulus window. **G)** Difference in peak Z-score for each stimulus between ACC→dlPAG and ACC broad excitatory recording groups. **H)** Difference in response decay for each stimulus between ACC→dlPAG and ACC broad excitatory recording groups. All error bars indicate SEM.

We found that L5p ACC→dlPAG neurons showed increased activity in response to all stimuli (**Figs. 4C, S11A**), but with varying response magnitudes (**Fig. 4D**). We observed that the larger the L5p ACC→dlPAG neural response, the longer the behavioral responses, regardless of whether the stimulus was noxious (**Fig. 4D**, R2 = 0.48, p < 0.001). We similarly observed striking aversive responses when recording from all pyramidal ACC neurons (**Fig. 4E,F**, R^2^ = 0.69, p < 0.001) and similar peak responses across stimuli (**Fig. 4G**, 2-way repeated measures ANOVA, p = 0.0516 with respect to virus). However, the L5p ACC→dlPAG neurons showed a significantly sharper decay than the general excitatory ACC population (**Fig. 4H**, 2-way repeated measures ANOVA, p = 0.0126, post-hoc comparisons of the AAVretro-GCaMP6s mice to the AAV-Camk2ɑ-GCaMP6s mice, Benjamini-Hochberg correction: miss: 0.5593, light hair: 0.4016, pin: 0.3231, Acetone: 0.5593, hot water: 0.1872, puff: 0.1872, n = 11 ACC→dlPAG, 4 ACC all-pyramidal).

Given our observation that ACC neural activity increases in response to all applied aversive stimuli, we next asked whether ACC activity reflects a conjunction of stimulus salience and valence rather than salience alone. In contrast to the increased neural activity observed during an aversive stimulus, we found a striking decrease in the activity of both L5p ACC→dlPAG and in the general ACC pyramidal neurons during free-foraging consumption of an appetitive stimulus (*i.e.,* chocolate, **Fig. S12G-J**). Further, we confirmed that this effect is not the result of foraging movement alone (**Fig. S12K-N**). Altogether, our data here show that ACC excitatory neurons respond broadly to salient stimuli and that L5p ACC→dlPAG neurons in particular exhibit more time-locked responses than the general population.

### L5p ACC→dlPAG neuron activity encodes heightened negative affect in a neuropathic state but does not strictly map to valence

Next, we asked how ACC neural dynamics evolve after the development of chronic neuropathic pain. In the same animals as in **Fig. 4**, we performed SNI and, after three weeks, retested the acetone and mechanical stimuli that correspond to the hallmarks of neuropathic hypersensitivity (**Fig. 5A,B**). L5p ACC→dlPAG neurons in injured mice responded similarly to those in uninjured mice, with more pronounced behavioral responses correlating with stronger calcium activity (**Fig. 5C,D, S11B,E,F**, R2 = 0.51, P < 0.001). However, the peak response of these neurons increased in response to all measured stimuli compared to the uninjured state (**Fig. 5E**, 2-way repeated measures ANOVA, p < 0.001 with respect to nerve injury state, post-hoc comparisons of acute to chronic pain, Benjamini-Hochberg correction for multiple comparisons: Miss: 0.00546, Light hair: 0.0194, Pin: 0.0194, Acetone: 0.0777). In contrast, general ACC pyramidal neurons showed no change in either the response magnitude or decay between the baseline and chronic pain states (R2 = 0.58, P < 0.001) (**Fig. 5F-H, S11H,K,L)**. The mean peak responses for L5p ACC→dlPAG neurons increased after SNI relative to the general ACC pyramidal neuron population (**Fig. 5I**, 2-way repeated measures ANOVA, p < 0.01 with respect to virus, post-hoc comparisons of ACC→dlPAG to ACC all-pyramidal, Benjamini-Hochberg correction: Miss: 0.1956, Light hair: 0.1956, Pin: 0.04724, Acetone: 0.1956), but the decay shape of the traces stayed the same (**Fig. 5J**, 2-way repeated measures ANOVA, p = 0.3181 with respect to virus). These data show that L5p ACC→dlPAG neurons exhibit a marked increase in response magnitude after the onset of chronic neuropathic pain, which is not present in the general ACC pyramidal neuron population.

**Figure 5.**
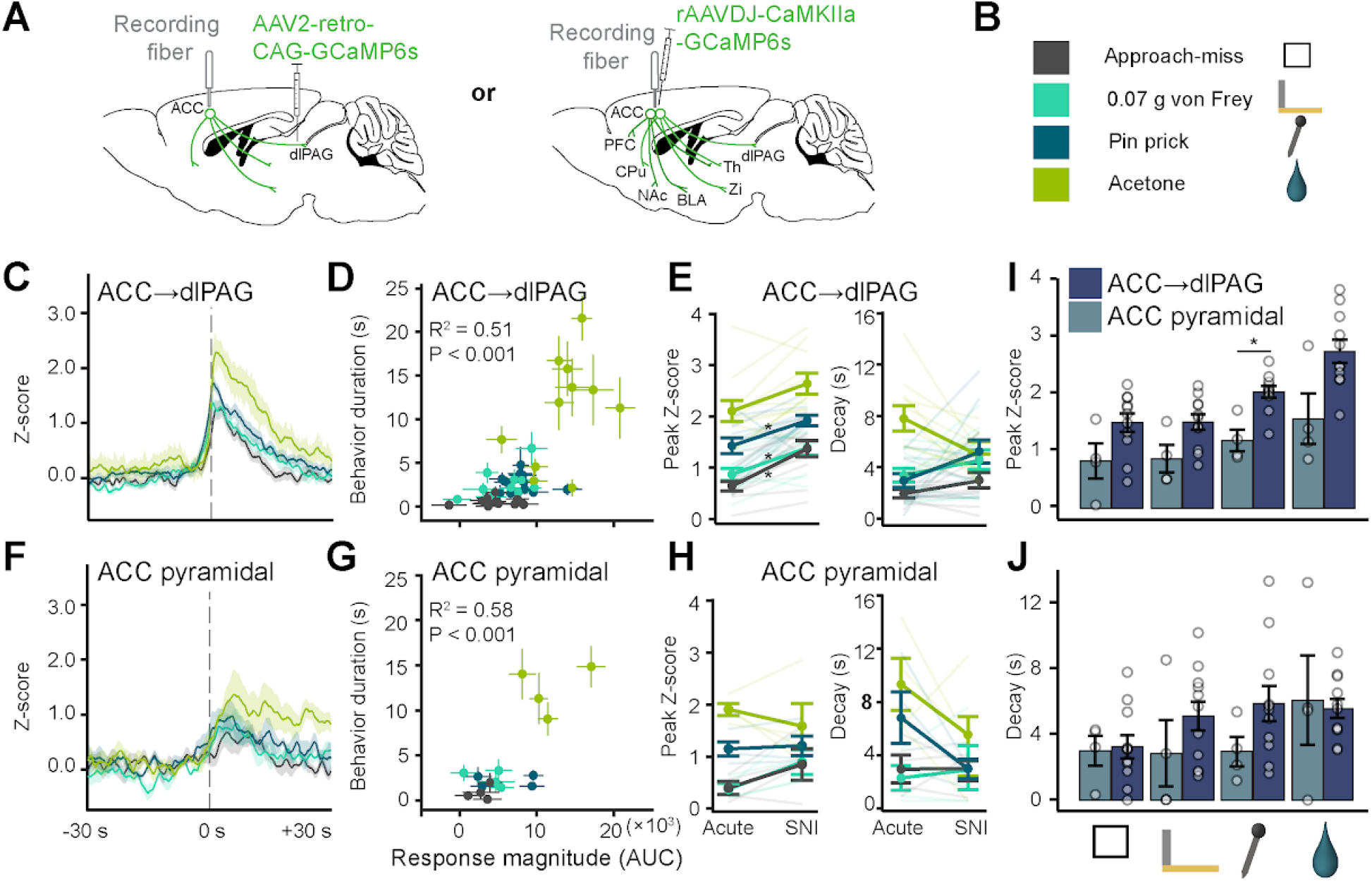
ACC→dlPAG neurons show potentiated activity to aversive stimuli in the chronic pain state. **A)** Virus injection schematic to selectively record activity of either ACC→dlPAG projection neurons (*Left*) or all ACC excitatory pyramidal neurons (*Right*). Injections and implants were unilateral. **B)** Description of stimuli with color code and icon legend. **C)** ACC→dlPAG mean fluorescence trace ± SEM during stimulus application three weeks following nerve injury. **D)** Pearson correlation of ACC→dlPAG response magnitude and behavior duration in the 60-second post-stimulus window. Error bars reflect SEM for individual trials on the respective axis. **E)** ACC→dlPAG peak responses (*Left*) and decay (*Right*) between acute and chronic pain. **F)** ACC broad excitatory population mean fluorescence trace ± SEM during stimulus application, three weeks following nerve injury. **G)** Pearson correlation of ACC broad excitatory response magnitude and duration of pain behavior in the 60-second post-stimulus window. Error bars reflect SEM for individual trials on the respective axis. **H)** ACC broad excitatory peak response (*Left*) and decay (*Right*) response values for the four tested stimuli between acute and chronic pain. **I)** Difference in peak Z-score for each stimulus between ACC→dlPAG and ACC broad excitatory recording groups after SNI. **J)** Difference in response decay for each stimulus between ACC→dlPAG and ACC broad excitatory recording groups. All error bars indicate SEM.

Finally, we used these continuous fiber photometry recordings to examine the relationship between behavior and ACC activity, in both the presence and absence of stimulus. To this end, we overlaid the epochs of recorded behaviors with the fluorescence signal recordings (**Fig. 6A,D**). After noxious stimulation of the hindpaw, we observed significant concurrence of neural activity and pain behaviors, but not grooming or rearing, for both L5p ACC→dlPAG and general ACC pyramidal neurons (**Fig. 6B,C,E,F**). Similarly, mice with SNI showed increased neural activity in both populations, concomitant with pain behaviors but not non-nociceptive behaviors (**Fig. S12A-D**).

**Figure 6.**
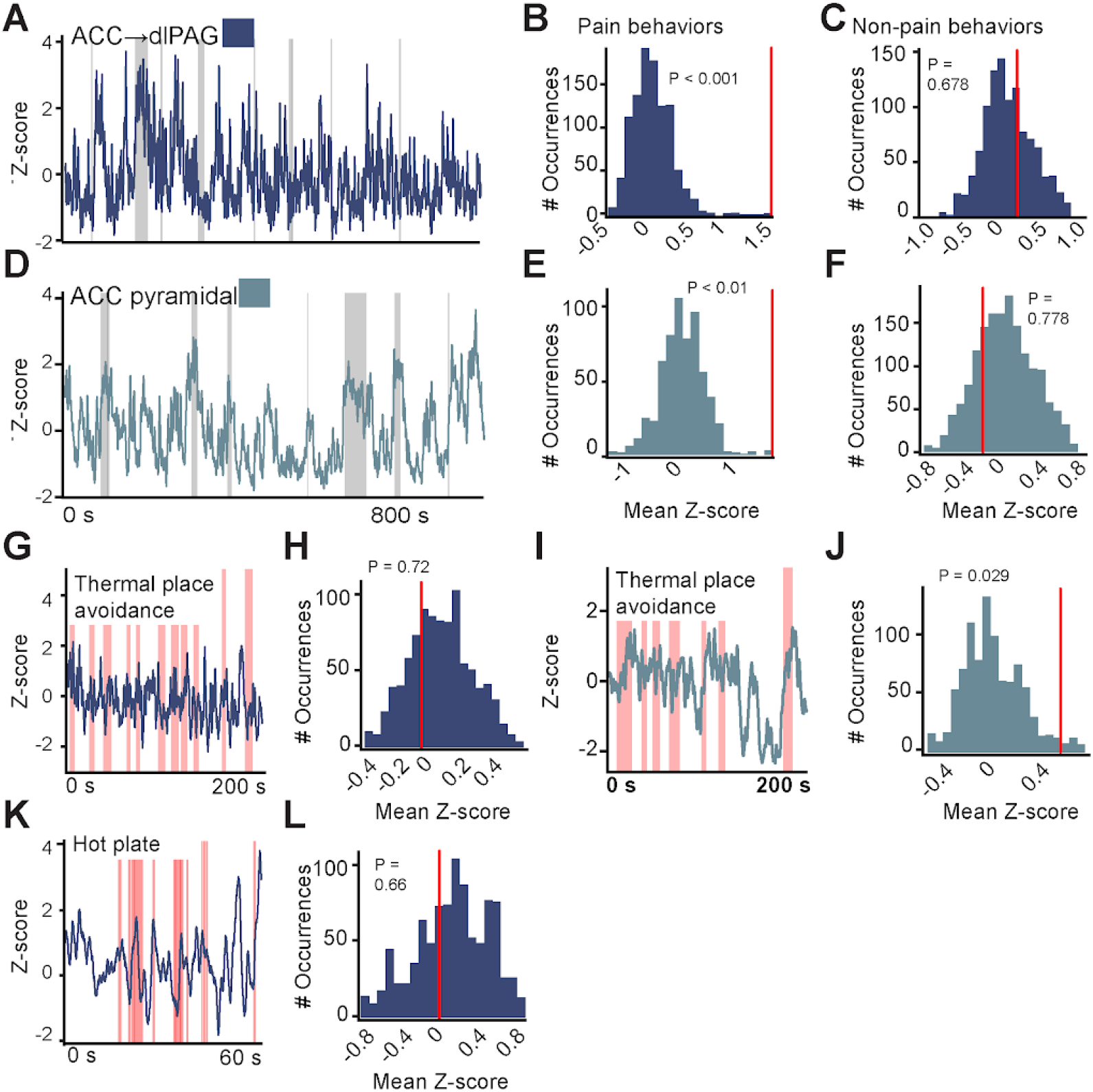
ACC pyramidal neurons increase activity during spontaneous pain behaviors. **A)** Representative trace from an ACC→dlPAG fiber photometry recording in an uninjured animal with transparent gray bars overlaying epochs of pain behaviors. **B)** Representative histogram from a permutation test (1000 shuffles), in which the timing of the pain behavioral epochs was linearly translated by random variation from the corresponding recorded trace in panel B. **C)** Representative histogram from a permutation test for non-pain behaviors from the trace in panel B. **D)** Representative trace from an ACC broad excitatory neuron fiber photometry recording in an uninjured animal with transparent gray bars overlaying epochs of pain behaviors. **E)** Representative histogram from a permutation test for pain behaviors in panel D. **F)** Representative histogram from a permutation test for non-pain behaviors from the trace in panel D. **G)** Thermal place avoidance representative trace from an ACC→dlPAG recording with transparent red bars overlaying hot side arena visits. **H)** Representative histogram from a permutation test for hot side visits from recording in panel G. **I)** Thermal place avoidance representative trace from an ACC broad excitatory recording with transparent red bars overlaying hot side arena visits. **J)** Representative histogram from a permutation test for hot side visits from recording in panel I. **K)** Hot plate representative trace from an ACC→dlPAG recording with transparent red bars overlaying pain behavioral epochs. **L)** Representative histogram from a permutation test for pain behavior epochs from recording in panel K.

While these ACC neurons respond to discrete noxious stimuli, when the noxious stimulus was persistently present, such as in the thermal place avoidance and hot plate tests, we found no correspondence between ACC neuronal activity and the animal’s presence on the noxious side or escape behaviors in the thermal place avoidance test (**Fig. 6G-J**), and no relationship between activity peaks and affective-motivational behaviors in the hot plate test (**Fig. 6K,L**). This result suggests that ACC activity does not directly reflect the beginning and end of pain affective-motivational behaviors themselves. Altogether, these data implicate ACC neural activity in the affective-motivational component of pain but not the sensory experience or reflexive responses.

## Discussion

These studies identify a subpopulation of nociceptive excitatory layer 5 ACC neurons that project to the dlPAG but not to the BLA or spinal cord dorsal horn, forming monosynaptic connections with primarily glutamatergic dlPAG neurons (**Fig. 1**). In both uninjured mice and mice with a sciatic nerve injury, chemogenetic inhibition of ACC→dlPAG neurons reduced affective-motivational pain behaviors, without affecting stimulus detection or withdrawal reflexes (**Figs. 2,3**). Calcium activity recordings in the ACC of freely behaving mice showed that ACC→dlPAG neurons are activated by both noxious and innocuous aversive somatosensory stimuli; however, the ACC→dlPAG population shows sharper tuning to the stimulus onset, whereas the global *Camk2*ɑ+ population of ACC neurons shows a broader time course of excitation and decay (**Fig. 4**). Furthermore, during nerve injury–induced chronic pain, the ACC→dlPAG population exhibits potentiated stimulus responses that are not observed in the global *Camk2*ɑ+ population (**Figs. 5,6**). Taken together, these results suggest that L5p ACC→dlPAG neurons contribute to encoding the affective dimension of pain without affecting sensitivity to noxious stimuli.

Although there is broad agreement that the ACC contributes to aversion and affective-motivational behaviors during pain experience, reports conflict as to whether the ACC also modulates nociceptive thresholds and withdrawal reflexes [(Calejesan et al., 2000; Gu et al., 2015; Koga et al., 2015; Li et al., 2010; Oswell et al., 2026; Santello and Nevian, 2015) *vs.* (Barthas et al., 2015; LaGraize et al., 2004; Lee et al., 2022)]. Theoretically, these inconsistent results could stem from distinct ACC neuron types that exclusively or preferentially influence pain affect, whereas others additionally modulate sensitivity to noxious stimuli or other ACC functions, such as empathy and fear. In support of this hypothesis, recent work has shown that projection-specific pathways from the ACC to the NAc and BLA separately affect the social transfer of pain and fear (Smith et al., 2021).

Our work similarly supports this theory. First, we found that L5p ACC→dlPAG projection neurons do not project or have particularly sparse projections to the spinal cord dorsal horn (Chen et al., 2018), RVM (*i.e.,* the node of the descending pain control system that is the downstream gateway to the dorsal horn (Heinricher et al., 1987) (Calejesan et al., 2000), or BLA (Smith et al., 2021), thus identifying another distinct pyramidal neuron subpopulation in ACC (**Figs. 1, S2**). Second, our behavioral data from chemogenetic inhibition of ACC→dlPAG provide additional evidence that pain affect and nociceptive threshold can change independently, suggesting that different brain circuits may modulate distinct dimensions of the pain experience (**Fig. 2**). This circuit-level dissociation of the sensory and affective components of pain presents an opportunity to create novel therapies that render noxious stimuli emotionally inert, eliminating pain suffering without disrupting the beneficial protective functions of noxious stimulus detection, such as withdrawal reflexes (Bushnell et al., 2013; Cox et al., 2006; De Ridder et al., 2021; Rainville et al., 1997).

The pattern of correlations between ACC response magnitude and aversion intensity suggests that the ACC processes an epiphenomenon of aversion rather than unpleasantness itself (**Figs. 4, 5**). Both our global ACC and ACC→dlPAG recordings show a positive trend in which response magnitude correlates with nocifensive behavior duration. However, we observed a stronger correlation in CamkIIɑ+ ACC than in ACC→dlPAG for response magnitude with aversion intensity, with behavioral duration serving as a proxy for intensity. This relationship, coupled with the sharper ACC→dlPAG neuron responses relative to Camk2ɑ+ neuron responses, suggests that it is not merely a random sample of *Camk2*ɑ+ ACC neurons and supports the theory that distinct anatomical circuits process different components of global ACC computations. The lower sensitivity of ACC→dlPAG neurons to aversion intensity suggests that this circuit does not encode the degree of unpleasantness. Additionally, during the transition from acute to chronic pain, the correlation between both Camk2ɑ+ and ACC→dlPAG neuronal activity and aversive intensity weakens. If encoding unpleasantness were the true function, we would expect the relationship between activity and behavior to be maintained during chronic pain, since the ceiling of aversive perception to the same stimuli is higher and the response magnitude is larger. Instead, ACC→dlPAG activity could be tied non-linearly to some threshold of inputs, indicating that something sufficiently threatening has occurred. This would explain the larger ACC→dlPAG responses relative to *Camk2*ɑ+ for stimuli such as the approach-miss and light hair, which are still perceptible events but are much less somatically intense than the other stimuli. Our data further show that even within a specific subpopulation of behaviorally necessary ACC neurons that are strongly responsive to noxious stimuli, the response profiles are not nociception-or salience-specific. This argues against the existence of specialized or dedicated “pain aversion” neurons in the ACC and instead suggests that pain activates ACC circuits that encompass pain but also encode broader features of aversive experience, such as autonomic responses (Wang et al., 2024). However, given that our recordings were at the population level, we cannot rule out subpopulations of ACC→dlPAG or ACC pyramidal neurons that are specifically encoded for pain.

Future work can address this with single-cell Ca+ recordings. One such study recently found that individual neurons in the ACC, including a subpopulation projecting to dlPAG (but notably not to the BLA), encode both observed fear aversion and shock-induced pain (Choi et al., 2025), supporting our conclusion that the ACC does not have pain-dedicated neurons.

The ACC→dlPAG neurons characterized in this study present a new opportunity to better understand how PAG circuits contribute to pain. The PAG can be spatially separated into columns with distinct behavioral and physiological functions (Bandler and Shipley, 1994; Vaughn et al., 2022). These PAG domains receive inputs from multiple other brain regions involved in sensory processing and internal representation, such as the cortex, hypothalamus, and amygdala (Keay and Bandler, 2015). Accordingly, neuronal populations within the PAG may act as effectors encoding specific behavioral outputs or discrete functional elements shared across multiple behaviors. Interestingly, recent single-cell spatial transcriptomics studies revealed that these regions appear transcriptionally distinct from one another (Vaughn et al., 2022). The ventral and dorsal columns are thought to be involved in passive *vs.* active defensive coping strategies, respectively (Keay and Bandler, 2015). The dorsal PAG has been proposed to regulate active coping strategies displayed during fear and aggression by modulating the cardiorespiratory system through its projection to the parabrachial nucleus (Evans et al., 2018; Falkner et al., 2020; Keay and Bandler, 2015; López-González et al., 2020). Yet, how the dlPAG specifically contributes to pain coping and defensive strategies remains unclear. Our data, combined with the known functional features of the dlPAG, suggest that ACC projections to this area may contribute to the emotional arousal associated with pain by modulating autonomic responses rather than sensory perception.

The diversity of threat-response behaviors mediated by the dlPAG (Behbehani, 1995; Keay and Bandler, 2015; F. M. C. V. Reis et al., 2021) suggests that behavioral output is not a simple relay function of inputs during an aversive experience. The relevant dlPAG-mediated behavior depends on the contextual demands of threat avoidance (Deng et al., 2016; Evans et al., 2018; Falconi-Sobrinho et al., 2024; Kim et al., 2013; F. M. Reis et al., 2021). In addition to accurately encoding defensive behaviors and escape, dlPAG neuron ensembles identify distance to the threat, predator speed, and angular offset from the threat (F. M. C. V. Reis et al., 2021). Our findings support a direct contribution of dlPAG activity to nocifensive pain behaviors, which adds to the repertoire of behaviors mediated by the dlPAG (Calejesan et al., 2000; Franklin et al., 2017; Vander Weele et al., 2018).

Finally, we observe selective potentiation in the ACC→dlPAG subpopulation between acute and chronic pain, but not the *Camk2*ɑ*+* pyramidal population. This demonstrates that not all neurons in the ACC contribute equally to chronic pain and further supports the idea that chronic pain potentiates acute pain circuits in the ACC. In complementary studies, PAG-projecting L5p and contralaterally projecting L5p ACC neurons exhibit distinct changes in their electrophysiological properties following CFA-induced chronic inflammatory pain (Franciosa et al., 2024), reinforcing our conclusion that discrete pyramidal populations contribute differentially to the development and expression of chronic pain.

A limitation of this study is that the behavioral and photometry experiments were conducted in male mice. Sex can influence pain-related behaviors and the neural mechanisms that support them; therefore, it remains to be determined whether the organization, recruitment, and functional contribution of the ACC→dlPAG pathway to acute and neuropathic pain-associated affective-motivational behaviors are conserved in females. Future studies that include both sexes will be important for establishing the generalizability of these findings.

In conclusion, this study provides evidence that ACC→dlPAG neuronal activity critically contributes to the aversive component of pain by regulating affective experience. These findings suggest that targeting the activity of ACC→dlPAG neurons could be an avenue towards better analgesic interventions to directly nullify the cause of pain’s distress.

## Materials and methods

### Key resources

| Reagent type | Designation | Source or reference | Identifiers | Additional information |
| --- | --- | --- | --- | --- |
| Strain, strain background (M. musculus) | C57BL/6J | Jackson Laboratory | Jax 000664 | Male, used for most experiments |
| Strain, strain background (M. musculus) | Vgat-Cre | Jackson Laboratory | Jax 016962 | Used for ex vivo electrophysiology |
| Strain, strain background (M. musculus) | Vglut2-Cre | Jackson Laboratory | Jax 016963 | Used for ex vivo electrophysiology |
| Virus | CAV-hSyn-Cre | Montpellier, France | — | $2 \times 10^{11}$ |
| Virus | AAV-retro-CAG-Cre | Kimberly Ritola (Janelia; UNC-NeuroTools) | — | $3.7 \times 10^{12}$ |
| Virus | rAAV5-Ef1a-DIO-hM4Di-mCherry | UNC Vector Core | — | $6.5 \times 10^{12}$ |
| Virus | rAAV5-hSyn-DIO-hM4Di-mCherry | Addgene | #44362 | $4.7 \times 10^{12}$ |
| Virus | rAAVDJ-Ef1a-DIO-mCherry | Stanford Vector Core | — | $8.5 \times 10^{12}$ |
| Virus | rAAV5-Camk2α-Chr2-EYFP | UNC Vector Core | — | $7.4 \times 10^{12}$ |
| Virus | rAAV5-Camk2α-EYFP | UNC Vector Core | — | $1.8 \times 10^{13}$ |
| Virus | rAAV5-hSyn-FLEX-GFP | UNC Vector Core | — | $8.7 \times 10^{12}$ |
| Virus | rAAVDJ-CamkIIα-GCaMP6s | Stanford Vector Core | GVVC-AAV-088 | $2.93 \times 10^{10}$ |
| Virus | rAAV2-retro-CAG-GCaMP6s | Kimberly Ritola (Janelia; UNC-NeuroTools) | — | $2.1 \times 10^{13}$ |
| Antibody | Anti-c-Fos (rabbit polyclonal) | Synaptic Systems | Cat# 226 003 | 1:5000, primary |
| Antibody | Donkey anti-rabbit Alexa Fluor 555 | Invitrogen (Fisherscientific) | # A31572 | Secondary, 1: not specified |
| Chemical compound, drug | Clozapine-N-oxide (CNO) | Tocris | #4936 | 10 mg/kg, i.p., in 0.9% saline |
| Chemical compound, drug | 4-hydroxytamoxifen | Sigma | T176 | 50 mg/kg in Kremofo EL vehicle, s.c. |
| Chemical compound, drug | Kremofo EL | Sigma | C5135 | Vehicle for 4-OHT |
| Software, algorithm | Ethovision XT 11.5 | Noldus | — | Behavioral tracking |
| Software, algorithm | LAS AF Lite | Leica Microsystems | — | Confocal image acquisition |
| Software, algorithm | pClamp9 | Molecular Devices | — | Electrophysiology acquisition |
| Software, algorithm | ImageJ | NIH | — | Manual stimulus timestamp scoring |
| Software, algorithm | Custom Python scripts | This paper | — | Photometry-video sync, analysis; available upon request |

### Animals

All animal procedures were approved by the Stanford University Administrative Panel on Laboratory Animal Care (APLAC, protocol #27378) and the University of North Carolina Institutional Animal Care and Use Committee (IACUC, protocol 24-052) and abided by the recommendations of the International Association for the Study of Pain (IASP). All animals, except for *in vitro* electrophysiology and one tracing experiment, were C57BL/6J male mice (JAX). *Vgat*-Cre and *Vglut2*-Cre mice were used for *in vitro* electrophysiology. Mice were housed 2–5 per cage and maintained on a 12-h light-dark cycle in a temperature-controlled environment with *ad libitum* access to food and water. All surgeries were performed on mice at least 8 weeks old. hM4Di-DREADD behavior experiments were performed 4–5 weeks after virus injections. In vitro electrophysiology was performed 3 weeks after virus injections.

### Viruses and surgery

All mice in which surgery was performed were anesthetized with isoflurane (5% for induction, 1.0– 2.0% thereafter) and placed in a stereotaxic frame, with their body temperature maintained using a heating pad. Paw color and breathing rate were monitored throughout the surgery. After the surgery, the skin on the head was sutured together (Ethicon sutures), and the mouse was given 1 mL of 0.9% saline.

#### Viruses used

CAV-hSyn-Cre (Montpellier, France), 2×10^11^ AAV-retro-CAG-Cre (Kimberly Ritola, initially Janelia Research Campus, then UNC-NeuroTools), 3.7×10^12^ rAAV5-Ef1a-DIO-hM4Di-mCherry (UNC Vector Core), 6.5×10^12^ rAAV5-hSyn-DIO-hM4Di-mCherry (Addgene), 4.7×10^12^ rAAVDJ-Ef1ɑ-DIO-mCherry (Stanford Vector Core), 8.5×10^12^ rAAV5-Camk2ɑ-ChR2-EYFP (UNC Vector Core), 7.4×10^12^ rAAV5-Camk2ɑ-EYFP (UNC Vector Core), 1.8×10^13^ rAAV5-hSyn-FLEx-GFP (UNC Vector Core), 8.7×10^12^ rAAVDJ-CamkIIɑ-GCaMP6s (Stanford Vector Core), 2.93×10^10^ rAAV2-retro-CAG-GCaMP6s (Kimberly Ritola, initially Janelia Research Campus, then UNC-NeuroTools), 2.1×10^13^*Syringe and needle.* For all injections, a WPI 10 uL syringe with a 25 G beveled needle was used. Each virus injection used a separate syringe to avoid contamination.

*Fluid delivery.* The stereotaxic arm was outfitted with a knob that, when manually turned once, delivers ∼30 nL of fluid at ∼1 nL per second.

#### Injection coordinates and procedure

For ACC: AP +1.25 from bregma, ML +-0.25, DV-1.0 from the needle bevel tip at the surface of the brain. The bevel was positioned to face medially, and the whole arm was angled at 4°. Approximately 200 nL of virus was delivered, followed by an additional 7 minutes of diffusion before the needle was slowly removed.

For PAG: AP-4.0 to-4.1 from bregma (depending on the age of the animal), ML +-0.8, DV-2.1 from the needle bevel tip at the surface of the brain. The bevel was positioned to face medially, and the whole arm was angled at 5°. Approximately 60 nL of virus was delivered, followed by an additional 7 minutes of diffusion before the needle was slowly removed.

#### Photometry fiber implant

Two weeks after GCaMP6s virus injection, a 0.48 NA optic fiber (Doric) was implanted into the ACC and affixed to the skull with cyanoacrylate superglue and dental cement (Stoelting).

### Pharmacology

10 mg/kg CNO (Tocris) was prepared in 0.9% saline. All CNO was administered intraperitoneally (i.p.) 60 min prior to behavioral testing.

### Histological preparation

Mice were transcardially perfused with 10% formalin in PBS. The brains and lumbar spinal cords were dissected from the mice, post-fixed in formalin for 24 hours, and then cryoprotected in 30% sucrose in PBS. Brains were frozen in O.C.T. (Sakura Finetek) and sectioned coronally at 40 μm using a cryostat (Leica Biosystems). For tissue with endogenous fluorescence requiring no immunohistochemistry, sections were washed in 0.1 M PB 3x at 10 minutes per wash, with 1:10,000 DAPI in the final wash to visualize cell nuclei.

### Fos immunohistochemistry

Tissue was washed with PBS 3x for 10 minutes, then blocked with PBS containing 5% normal donkey serum and 0.3% Triton X-100 for 1 hour at 23°C. The sections were then incubated with primary antibodies (Synaptic Systems c-Fos rabbit polyclonal, Cat. No. 226 003, at 1:5,000) at 4°C, overnight. After extensive washing with PBS containing 1% normal donkey serum and 0.3% Triton X-100, sections were incubated with a secondary antibody conjugated to Alexa Fluor 555 for 2 hours at 23°C. The final wash included 1:10,000 DAPI to visualize cell nuclei.

All images were collected at 20x with a Leica TCS SP5II confocal microscope with LAS AF Lite software (Leica Microsystems).

### Ex vivo slice electrophysiology

*Vgat*-Cre and *Vglut2*-Cre mice were prepared 3 weeks prior to electrophysiology testing with two virus injections: AAV5-Camk2a-ChR2-EYFP injected into the ACC and AAVDJ-Ef1a-DIO-mCherry injected into the PAG to visualize Cre+ cells. P63-70 *Vglut2*-Cre or *Vgat*-Cre mice were anesthetized with isoflurane and decapitated. The brain was rapidly removed and placed in oxygenated ice-cold dissection solution (in mM: 95 NaCl, 2.5 KCl, 1.25 NaH_2_PO_4_, 26 NaHCO_3_, 50 sucrose, 25 glucose, 6 MgCl_2_, 1.5 CaCl_2_, and 1 kynurenic acid, pH 7.4, 320 mOsm). The brain region containing the PAG was isolated and coronally sliced at 200 μm thick using a vibrating microtome (Leica, USA). Slices were incubated in oxygenated recovery solution (in mM: 125 NaCl, 2.5 KCl, 1.25 NaH_2_PO_4_, 26 NaHCO_3_, 25 glucose, 6 MgCl_2_, and 1.5 CaCl_2_, pH 7.4, 320 mOsm) at 35°C for 1 hour. Patch-clamp recordings in the whole-cell configuration were performed at RT on mCherry+ cells in the dorsal PAG, visualized with an Olympus BX51WI or Zeiss Axioskop microscope fitted with Nomarski optics and connected to a camera (Dage-MTI, USA). Slices were perfused at 1-2 mL/min with recording solution (recovery solution containing 1 mM MgCl_2_ and 2 mM CaCl_2_). Recordings were performed in voltage-clamp mode at a holding potential of-70 mV. Thick-walled borosilicate pipettes, having a resistance of 3-5 MOhm, were filled with internal solution (in mM: 120 K-methyl-sulfonate, 10 NaCl, 10 EGTA, 1 CaCl_2_, 10 HEPES, 0.5 NaGTP, 5 MgATP, pH adjusted to 7.2 with KOH, osmolarity adjusted to 305 with sucrose). Data were acquired using a Multiclamp 700A amplifier and pClamp9 software (Molecular Devices, USA). Sampling rate was 10 kHz and data were filtered at 2 kHz. EPSCs were evoked with blue light generated by an LED Transmitted Light Source (Lambda TLED, Sutter) at 5 Hz with a 10 ms pulse width. PAG cells were considered to receive monosynaptic inputs if EPSCs reliably followed this pulse train.

### Spared nerve injury (chronic neuropathic pain)

The left hind leg was shaved and cleaned with ethanol. An incision was made in the skin, and the muscles retracted to expose where the sciatic nerve trifurcates as the common peroneal, tibial, and sural nerves. The common peroneal and sural nerves were transected. The muscle was closed with 5-0 sutures (Ethicon), and the skin wound was closed with tissue adhesive (Vetbond).

### Behavioral testing

#### Pre-test habituation

All mice were habituated to individual containers for 3 days prior to any behavioral testing. Containers were approximately 12 x 10 cm with air holes in the lid and 1 cm of bedding material at the bottom. Mice were not left in their individual containers for longer than 6 hours at a time.

#### Thermal place preference test

Two hot plates (BioSeb) were joined together to form a continuous surface with a <1 cm gap between them. A 12.5 x 25 cm white opaque plastic enclosure was placed over the plates, with equal area given to each side. One hot plate was set to 45°C (hot), and the other to 30°C (neutral). The hot and neutral plates remained the same for all testing. All mice were administered CNO 60 minutes prior to testing and were individually sequestered thereafter. Mice were lowered into the arena by their tails into an opaque tube placed on the neutral side and allowed to settle for 2-3 seconds. The tube was lifted, and the test session formally began. Mice were allowed to freely explore the arena for 5 minutes. The arena and enclosure were cleaned with 70% ethanol between each trial. Behavioral tracking and scoring: for hM4Di experiments, behavior was scored in real time using an overhead camera and Ethovision XT 11.5 (Noldus). The center of the animal’s body was tracked to determine its location. For photometry experiments, behavior was manually scored offline for timestamps of behavioral state changes.

#### Cold place preference test

The same conditions as in the thermal place preference test were used, except that the plate temperatures were set to 30°C and 15°C. A separate arena enclosure was used in the thermal place preference test, with a different material texture, to prevent a display of contextual learning from any previous testing. The dimensions were 10 x 30 cm.

#### Modified hot plate test

To evaluate affective-motivational responses to a sustained, inescapable noxious thermal stimulus, mice were placed on a 50°C hot plate for 60 s. A video camera (Sony Handycam HDR-CX220) positioned on the side, level with the hot plate floor, was used to capture the mice’s movement. This allows us to analyze high-speed videos, blinded to plate temperature or treatment, to quantify the total time spent engaging in various behaviors.

#### Cold plate test

The same conditions as the hot plate test were used, except the plate was set to 10°C. Mice were left on the plate for 60 seconds.

#### Classification of mouse behaviors into reflexive and affective-motivational nociceptive responses

In response to cutaneous noxious stimuli, mice can display an array of behaviors. Withdrawal reflexes are characterized by a rapid retraction, digit splaying, or flicking of the affected paw. Affective-motivational responses include coordinated behaviors such as paw-grabbing, licking, or biting, prolonged lifting and “guarding” of the paw away from the stimulus source, and jumping, rearing, or hyper-locomotion. Paw withdrawal reflexes are a product of spinal and brainstem reflex circuits, while affective-motivational behaviors require brain processing. The time the animal spends engaged in these affective-motivational behaviors reflects the degree to which it perceives the stimulus to be aversive and is sufficiently motivated to reduce the aversive nature of the stimulus (by licking the tissue or guarding against the stimulus), or to escape the stimulus source. By contrast, withdrawal reflexes reflect the degree to which sensory afferents recruit withdrawal reflexes and have classically been used in studies of hypersensitivity. For the hot plate test, behavior from both hind paws was scored. For the cold plate test, only behavior from the left (injured) hindpaw was scored.

#### Punctate stimuli tests (hM4Di experiments)

Mice were acclimated for 30–60 min in the testing environment within custom red plastic cylinders (4” D) on a raised metal mesh platform (24” H). All stimuli were applied from beneath the mesh. With the exception of up-down von Frey tests, stimuli were applied at 60-second intervals (longer if the animal was too active). The behaviors were scored in two categories: 1) the initial reaction of whether the animal withdrew its paw at the application of the stimulus, and 2) the post-stimulus response in the following 60 seconds. Nocifensive (as described above) and escape behaviors were scored positively, and are taken to indicate the degree of ongoing supraspinal affective-motivational processing elicited by the stimulus.

### Targeted Recombination in Active Populations (TRAP) of ACC neural ensembles

For all TRAP procedures, stereotaxic unilateral injections of viral reagents occurred 3–5 weeks prior to TRAP. We habituated mice to a first testing room (room A) for three consecutive days. Execution of all TRAP procedures occurred in Room A. During these habituation days, no nociceptive stimuli were delivered, and no baseline thresholds were measured (i.e., mice were naïve to pain experience before the TRAP procedure). In room A, we placed individual mice in red plastic cylinders (10.16-cm D) with red lids on a raised, perforated, flat metal platform (60.96-cm H). The male experimenter’s lab coat was present in the testing room for the first 30 min of acclimation, and then the experimenter entered the room for the final 30 min of habituation; this was done to mitigate potential alterations to the animal’s stress and endogenous antinociception levels. To execute the TRAP procedure, we placed mice in their habituated cylinder for 60 min, and then a 25G sharp pin was applied to the central-lateral plantar pad of the left hindpaw (tibial-sural nerve paw innervation territory), once every 30 s over 10 min. Following the pin stimulations, the mice remained in the cylinder for an additional 60 min before injection of 4-hydroxytamoxifen (Sigma T176; 50 mg/kg in ∼0.25 mL of vehicle [Kremofor EL: Sigma C5135]; subcutaneous). After the injection, the mice remained in the cylinder for an additional 2 hrs to match the temporal profile for c-FOS expression, at which time the mice were returned to the home cage (Note: an immediate return to the home cage following the pin stimulations was considered, but ultimately avoided as potential safety-related neural activity could occur and thus TRAP neurons of putative positive valence in addition to the nociceptive ensemble). After completing all experiments, we perfused the mice and dissected the brains to verify viral expression and quantify the resulting anatomy. We excluded mice with off-target viral expression in the anterior cingulate cortex from the behavioral analysis.

### Testing protocol for photometry behavior experiments

#### Testing in uninjured mice

The total testing protocol was spread out over two days. On day 1, mice were acclimated for 10 minutes in the testing environment within custom red plastic cylinders (4” D) on a raised metal mesh platform (24” H). A 5-minute baseline session was recorded with no stimuli, followed by a 5-minute break, followed by three “blocks” with a 5-minute break in between each. Each block consisted of 6 stimuli applied 3 times in a pseudo-randomized order for a total of 18 stimuli. The stimuli used were the hot-water drop, acetone drop, pinprick, 0.07 g von Frey hair, air puff, and approach-miss. Each break involved turning off the LEDs. After the third block and final 5-minute break, a final 5-minute baseline was recorded with no stimuli. On day 2, a similar procedure was followed but with two blocks instead of three. After the second block and following break, mice were exposed to 5 minutes in the open field, followed by 10-15 minutes of chocolate foraging, followed by a 5-minute thermal place preference test, all interleaved with 5-minute breaks.

#### Testing in mice with chronic pain

The total testing protocol was conducted in a single day, three weeks after the SNI surgery. Mice were acclimated for 10 minutes in the testing environment within custom red plastic cylinders (4” D) on a raised metal mesh platform (24” H). Von Frey up-down thresholds were measured for the day 21 time point, followed by a 5-minute break. A 5-minute baseline session was recorded with no stimuli, followed by a 5-minute break, followed by three “blocks” with a 5-minute break in between each. Each block consisted of 4 stimuli, applied 5 times in a pseudo-randomized order, for a total of 18 stimuli. The stimuli used were the acetone drop, pin prick, 0.07 g von Frey hair, and approach-miss. After the final block and break, a second baseline was recorded with no stimuli.

#### Punctate stimuli tests (photometry)

All tests were performed in custom red plastic cylinders (4” D) on a raised metal mesh platform (24” H). All stimuli were applied from beneath the mesh. Stimuli were applied at 60-second intervals for uninjured testing and 120-second intervals for chronic pain testing.

#### Mechanical sensitivity

To evaluate mechanical reflexive hypersensitivity, we used a logarithmically increasing set of 8 von Frey filaments (Stoelting), ranging in grams from 0.007 to 6.0 g. These were applied perpendicularly to the plantar hind paw with sufficient force to cause a slight bending of the filament. A positive response was characterized as a rapid withdrawal of the paw away from the stimulus filament within 2 seconds. Using the up-down statistical method, 50% withdrawal mechanical threshold scores were calculated for each mouse and then averaged across the experimental groups. For punctate-stimulus experiments, only 0.4 or 0.07 g of hair was used. For uninjured animals, the von Frey hairs were applied to the center of the left hindpaw plantar surface. For nerve-injured animals, they were applied to the outer “heel” of the paw, in the specific hypersensitive area of each individual mouse.

*Pin prick.* A 29.5 G insulin needle was used to poke the center of the left hindpaw plantar surface.

*Hot water drop.* A 50 uL drop of approximately 55°C hot water was applied to the left hindpaw by forming a surface tension bubble of liquid at the tip of a 100 uL syringe with no needle.

*Acetone drop.* A 50 uL drop of acetone was applied to the left hindpaw by forming a surface tension bubble of liquid at the tip of a 100 uL syringe with no needle.

*Air puff.* A rapid puff of air was applied to the hindpaw by rapidly squeezing a soft rubber puffer (Giottos rocket blaster).

*Approach-miss.* For the photometry experiments, to determine whether calcium transients were responses to the stimuli themselves or merely the anticipation of them, an “approach-miss” technique was used. Either the pin or the acetone syringe was raised near the animal’s hindpaw to recapitulate the act of stimulus application, but the stimulus was not applied. Instead, the tip of the stimulus device was held <1 cm away from the hindpaw for 3-4 seconds, then removed.

*Open field.* For hM4Di experiments, mice were placed in a 60-cm-diameter white cylinder for 20 minutes. Behavioral tracking and scoring were conducted using an overhead camera and Ethovision XT 11.5 (Noldus). For photometry experiments, a 35 x 40 cm rectangular white plastic arena was used.

*Chocolate.* In the photometry open field arena, 4-5 ∼80 mg chunks of milk chocolate were randomly scattered across the arena. To acclimatize animals to the food, 8-10 similarly sized pieces of chocolate were placed in their home cage for 3 days prior to the experiment. There was no food or water restriction. The chocolate exploration session was terminated after 15 minutes, regardless of how many chocolate pieces the mice consumed.

*Formalin.* 10 µl of 5% formalin in PBS was injected subcutaneously into the plantar surface of the left hindpaw. A video camera (Sony Handycam HDR-CX220) placed underneath the wire mesh recorded their behavior and movement for 60 minutes after injection. Mice were sacrificed and perfused 90 minutes after injection.

### Fiber photometry

Fluorescence excitation was provided by two LEDs at 488 nm and 405 nm (Thorlabs). The 488 LED was used to excite GCaMP6s, and the 405 LED served as a concomitant control. A real-time signal processor (RX8, Tucker-Davis Technologies) running custom software sinusoidally modulated each LED’s output (488 at 230 Hz and 405 at 330 Hz) through a Doric fluorescence minicube, and simultaneously demodulated the two output signals from the output of the photodetector. The two output signals were projected onto a photodetector (model 2151 femtowatt photoreceiver, Doric) and collected at a sampling frequency of 380 Hz.

The d*F*/*F* signal was calculated by fitting a linear least-squares fit between the 405 and 488 time series. All the time points from the 405 time series were passed through the least-squares fit function to generate a fitted 405 signal. Change in fluorescence (d*F*) was calculated as the 488 nm signal minus the fitted 405 nm signal, and d*F*/*F* was calculated by dividing each point in d*F* by the 405 nm fit at that time. For each recording session, all traces were smoothed with a 1000-ms sliding window average and Z-scored to account for changes between animals related to GCaMP6s expression levels or fiber tip placement.

Video was recorded with a Logitech C922x webcam at 30 fps. A custom-written Python script synchronized the onset of photometry with that of video recording. All timestamps for stimulus application were scored by hand using ImageJ.

### Sample size, randomization, and blinding

Group sizes were consistent with standard practice and power analyses conducted for animal use planning (alpha = 0.05, two-tailed; power = 80%; Lenth’s sample size calculator) based on variability and effect sizes observed in prior studies using similar behavioral, chemogenetic, and photometry paradigms. Animals were pseudorandomly assigned to treatment groups (e.g., saline vs. CNO, viral construct). Somatosensory stimuli (von Frey filament, pinprick, hot water, acetone) were presented in a pseudorandomized order across trials to prevent order effects. Experimenters were blinded to treatment group during behavioral scoring and, where applicable, during data analysis.

### Replicates

Unless otherwise noted, n refers to the number of biologically independent animals per group. For *ex vivo* slice electrophysiology experiments, n indicates the number of individual neurons recorded (technical replicates); the number of animals contributing cells is reported in the corresponding figure legends.

## Statistics

Statistical analyses were performed in Python and GraphPad Prism. Two-group comparisons were assessed with Welch’s unpaired t-test to account for unequal variances between groups. For repeated measurements across multiple time points or stimuli within the same animals, we used two-way repeated-measures ANOVA, testing main effects of treatment (or virus/nerve-injury state) and time (or stimulus), along with their interaction. For comparisons across more than two related stimulus conditions within a single population, we used one-way repeated-measures ANOVA. Where an ANOVA revealed a significant main effect, we performed post hoc pairwise comparisons (paired or unpaired t-tests, as appropriate to the comparison) with correction for multiple comparisons using the Benjamini-Hochberg procedure (false discovery rate). Proportions of monosynaptically connected neurons across genetically defined populations were compared using a 2-proportion Z-test. Associations between fiber photometry response magnitude and behavioral output were assessed by Pearson correlation. To determine whether neural activity was temporally coupled to specific behavioral epochs beyond chance, we used a permutation test in which behavioral epoch timing was randomly shuffled relative to the fluorescence trace over 1000 iterations to generate a null distribution, against which the true co-occurrence was compared. Significance was set at α = 0.05 (P<0.05, P<0.01, P<0.001, ****P<0.0001). All error bars represent SEM.

## Data and code availability

All data underlying the figures, including processed fiber photometry, behavioral, histological, and electrophysiological data, and the custom Python code used for analysis and figure generation, are available from the corresponding author upon reasonable request.

## Acknowledgments

This project was supported by the National Institutes of Health grant R01NS106301 (G.S.), the New York Stem Cell Foundation (G.S.), and the NSF Graduate Research Fellowships Program (GRFP) (J.K.). We thank J. Blair for manuscript editing.

## Author Contributions

J.K., G.C., and G.S. designed experiments and data analyses. J.K. performed and analyzed behavioral experiments. J.K. and W.M. performed and analyzed histological experiments. A.F. performed and analyzed electrophysiological experiments. E.K. and K.R. supplied viruses. J.K., W.M., and N.M.L. prepared the figures. J.K., A.F., N.M.L., and G.S. wrote the manuscript.

---

**Supplementary Figure 1.**
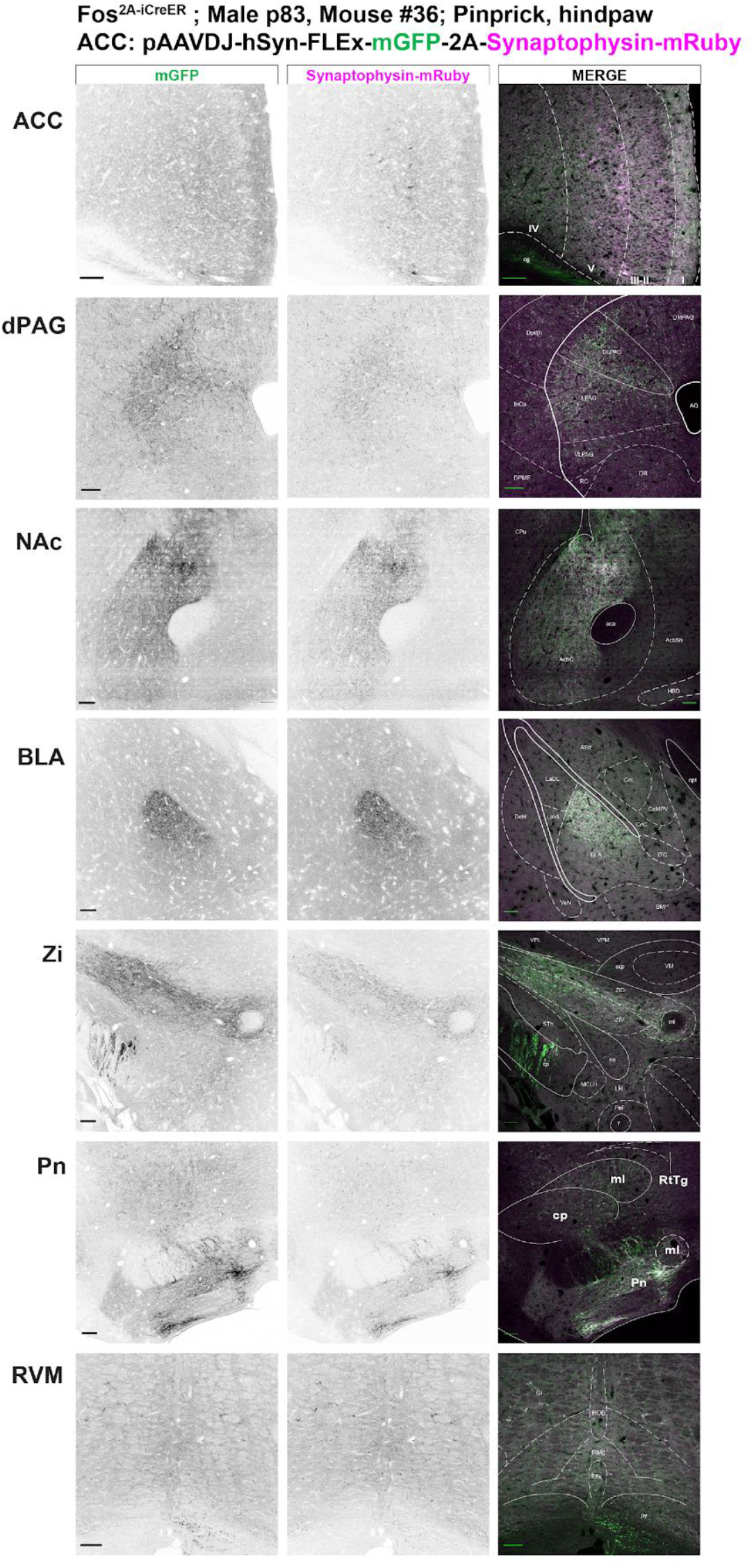
Nociceptive ACC neuron outputs. Example coronal images of mGFP and Synaptophysin-mRuby (inverse grayscale) and merge for the ACC injection site in TRAP2 mice with hindpaw pinprick and the outputs of ACC nociceptive neurons in the dPAG, NAc, BLA, Zi, Pn, and RVM. All scale bars = 100 µm.

**Supplementary Figure 2.**
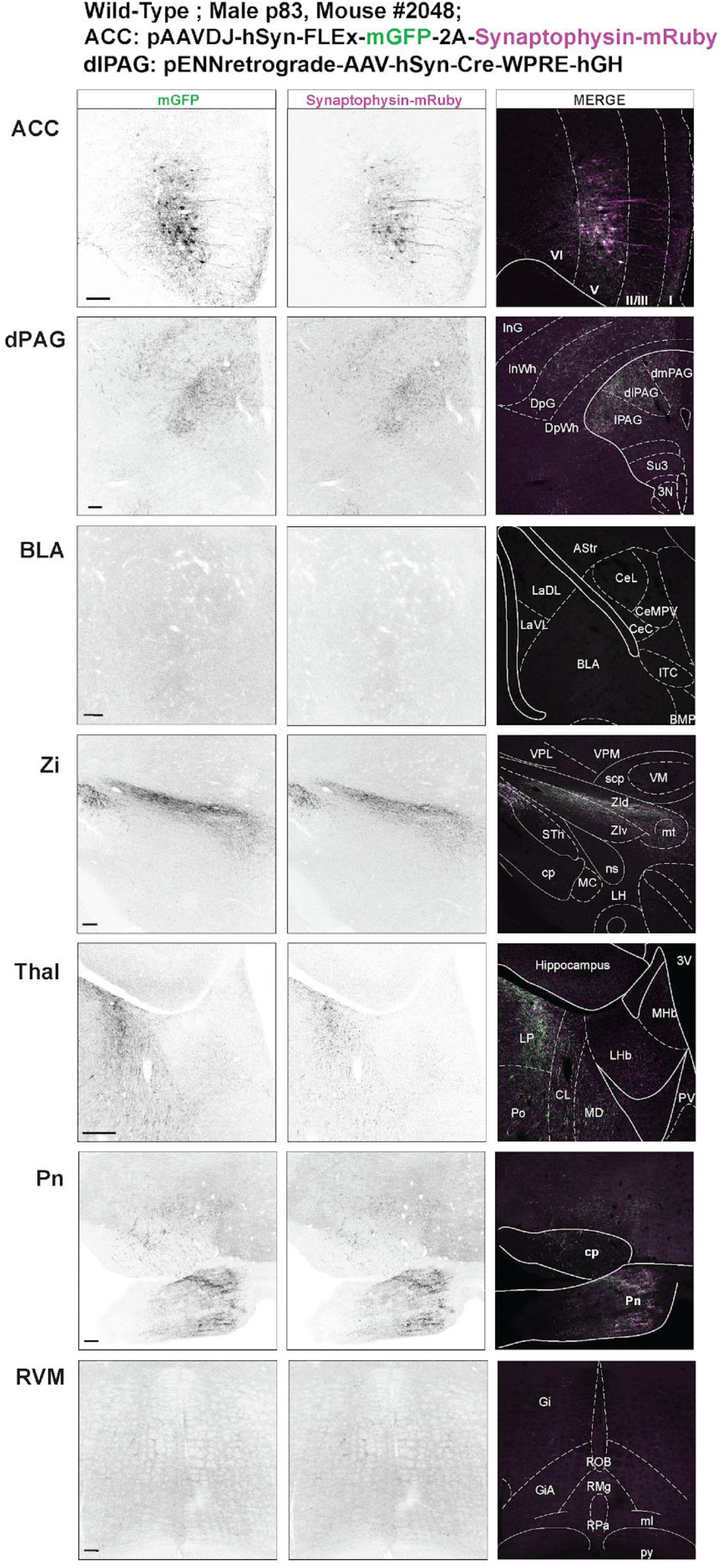
ACC→dlPAG collateral tracing. Example coronal images of mGFP and Synaptophysin-mRuby (inverse grayscale) and merge for the ACC and dPAG injection sites in wild type mice and the presence of absence of collaterals in the BLA, Zi, Thal, Pn, and RVM. All scale bars = 100 µm.

**Supplementary Figure 3.**
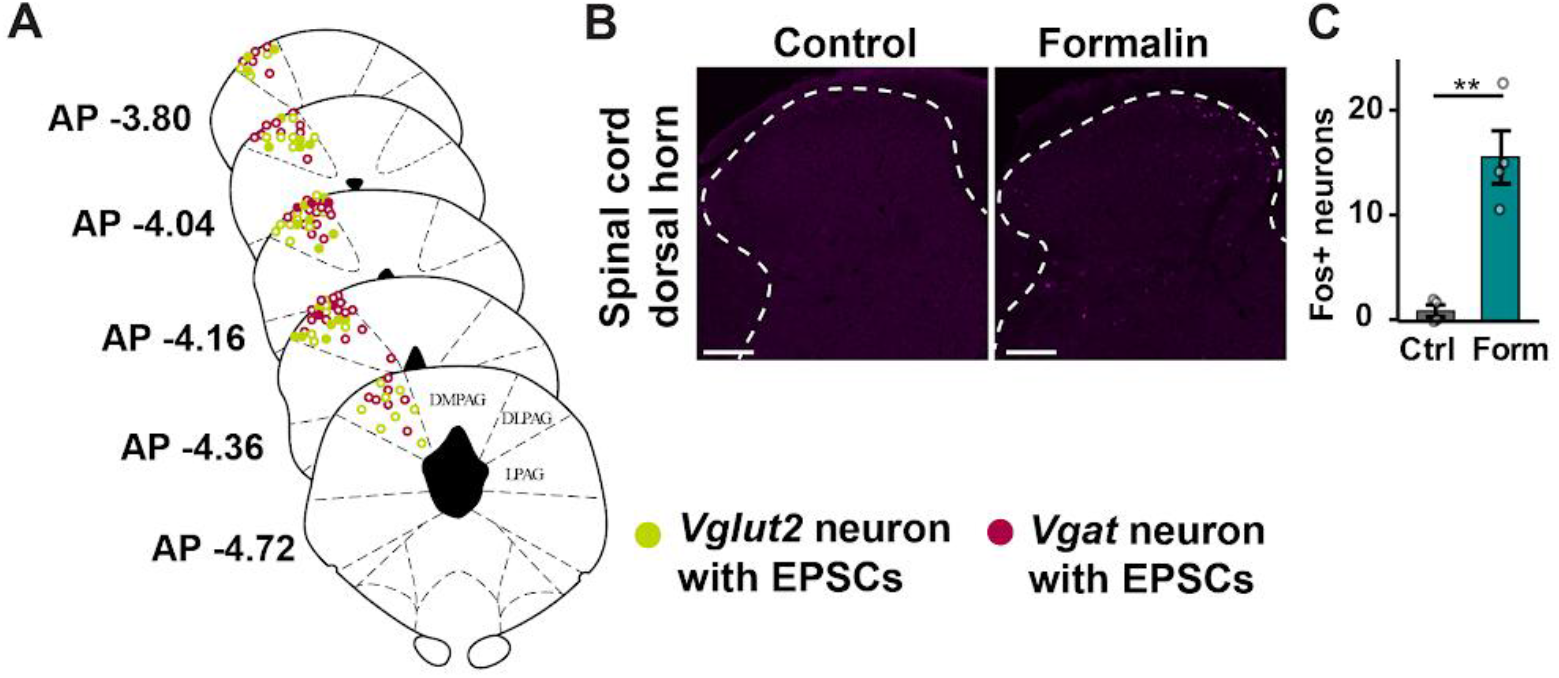
Extended data for patch locations and formal Fos quantification in. Figure 1**. A)** Location map of patch recordings. Filled circles are neurons with light-responsive EPSCs, open circles are neurons with no responses. **B)** Fos immunohistochemistry in the spinal cord dorsal horn after hindpaw formalin injection. **C)** Mean FOS counts per animal in the spinal cord. **p < 0.01. All scale bars = 100 µm. All error bars indicate SEM.

**Supplementary Figure 4.**
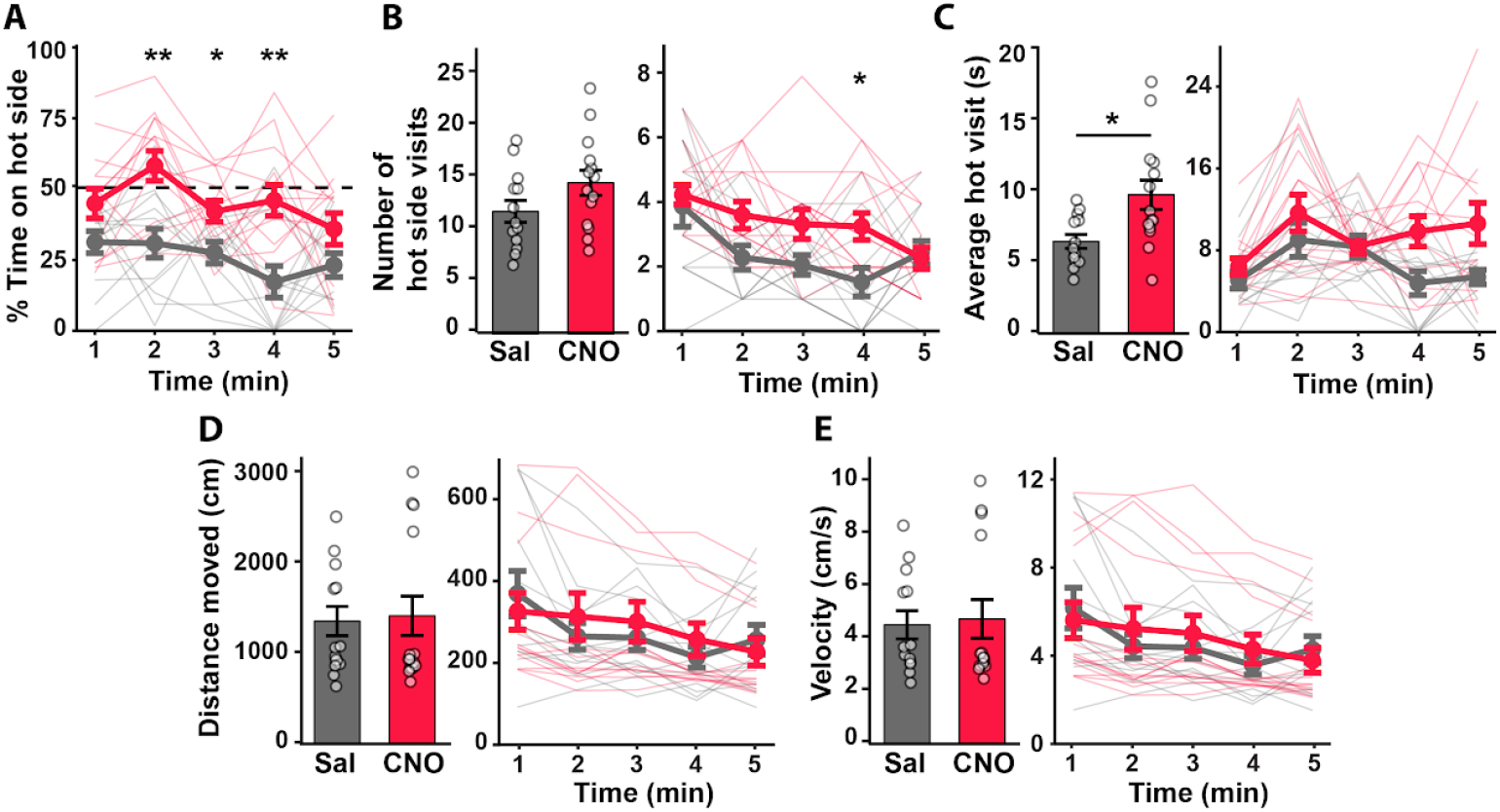
hM4Di ACC→dlPAG inhibition during thermal place avoidance, extended data. **A)** Percentage of hot zone time binned in 1-minute sequential segments. **B)** Number of hot side entries for the whole test session (Left) and number of hot side entries minute by minute (Right). **C)** Average duration of hot side visits (total time on hot side divided by number of visits) for the whole test session (Left) and average hot side visit duration minute by minute (Right). **D)** Total path distance traveled (Left) and total distance minute by minute (Right). **E)** Average velocity (Left) and average velocity minute by minute (Right). *p < 0.05, **p < 0.01. All error bars indicate SEM.

**Supplementary Figure 5.**
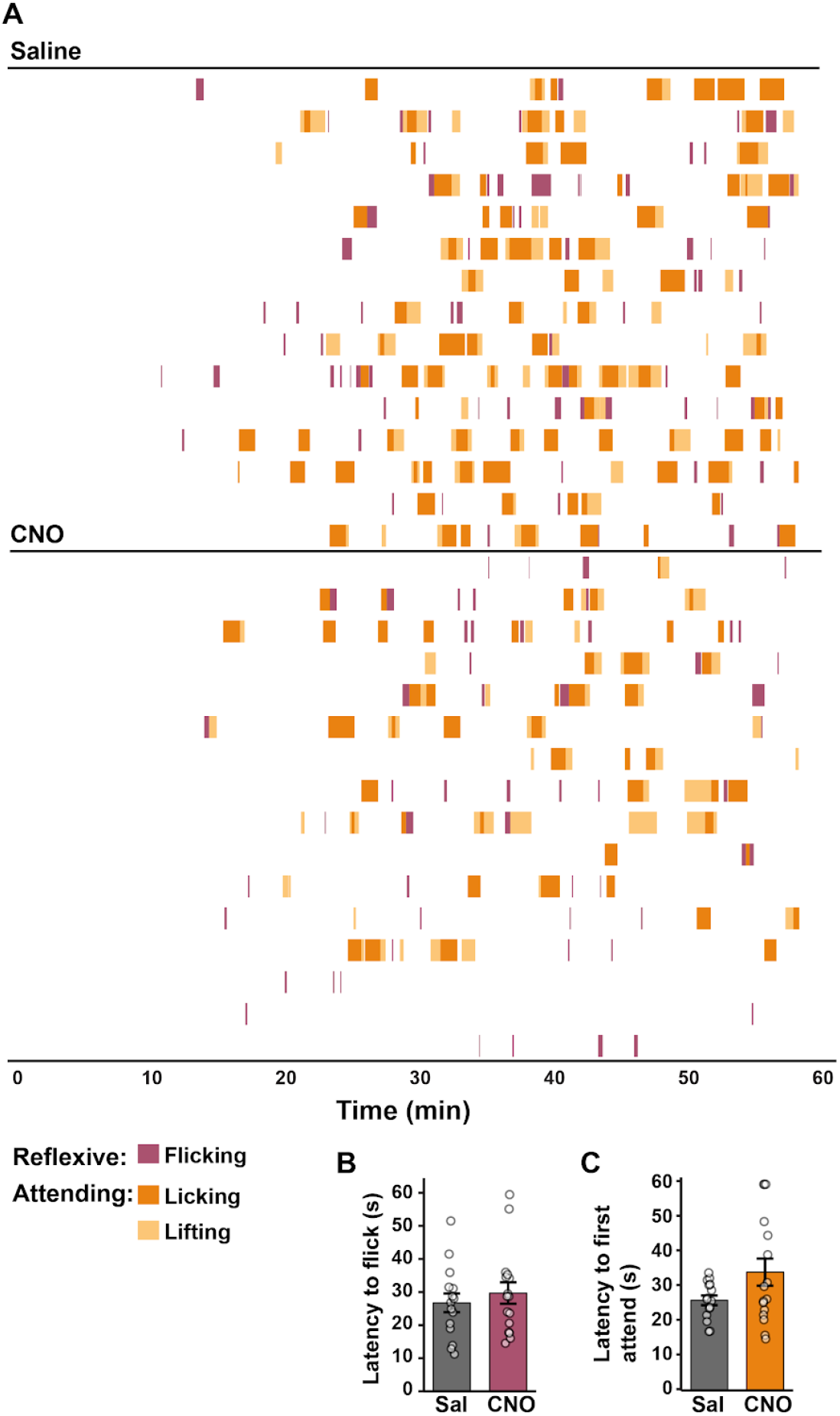
hM4Di ACC→dlPAG inhibition hot plate extended data. **A)**Behavior raster plots for all mice in each test group. Each row represents the behavior of one mouse over the course of 60 seconds on the hot plate. **B)** Latency to the first flick. **C)** Latency to the first lift or lick. All error bars indicate SEM.

**Supplementary Figure 6.**
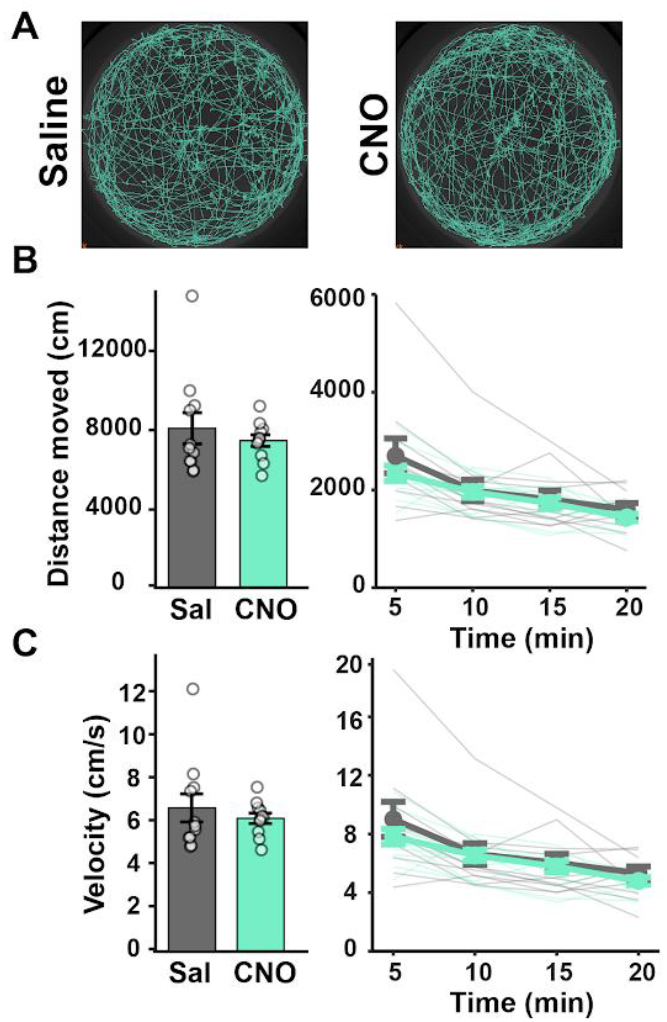
hM4Di ACC→dlPAG inhibition does not alter locomotion. **A)** Representative path plots for mice in each test group in the open field. **B)** Total distance moved over the whole session (*Left*), and distance moved binned into 5-minute segments (*Right*). **C)** Average velocity (*Left*) over the whole session and average velocity binned into 5-minute segments (*Right*). All error bars indicate SEM.

**Supplementary Figure 7.**
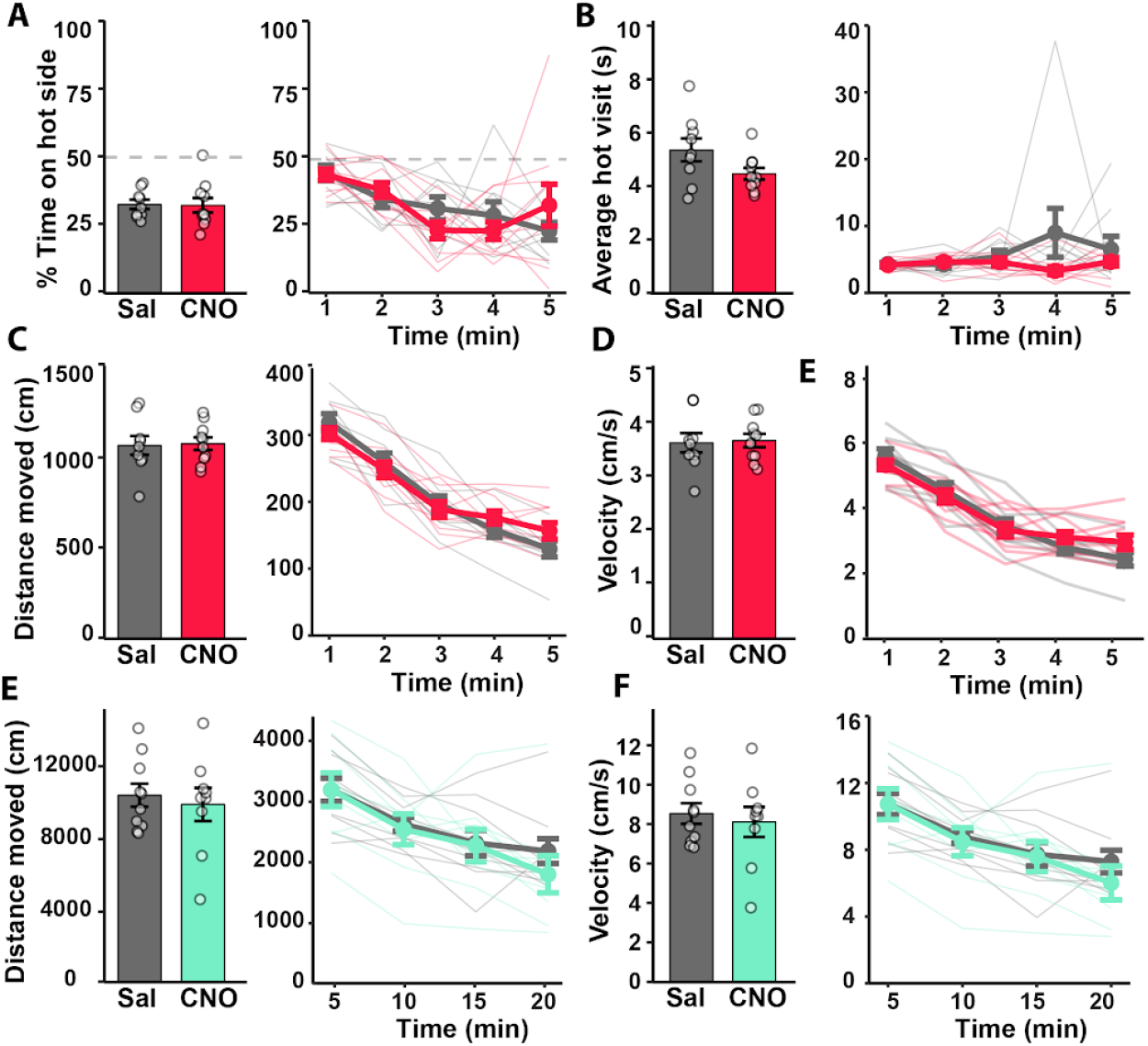
CNO alone in uninjected mice does not mediate pain, locomotion, or behavior. **A)** Percentage of time the mice spent on the hot side for the whole session (*Left*) and percentage of time the mice spent on the hot side, minute by minute (*Right*). **B)** Average duration of hot side visits for the whole test session (*Left*) and average hot side visit duration minute by minute (*Right*). **C)** Total distance moved over the whole thermal place avoidance session (*Left*), and distance moved binned into 1-minute segments (*Right*). **D)** Average velocity over the whole thermal place avoidance session (*Left*) and average velocity binned into 1-minute segments (*Right*). **E)** Total distance moved over the whole open field session (*Left*), and distance moved binned into 5-minute segments (*Right*). **F)** Average velocity over the whole open field session (*Left*) and average velocity binned into 5-minute segments (*Right*). All error bars indicate SEM.

**Supplementary Figure 8.**
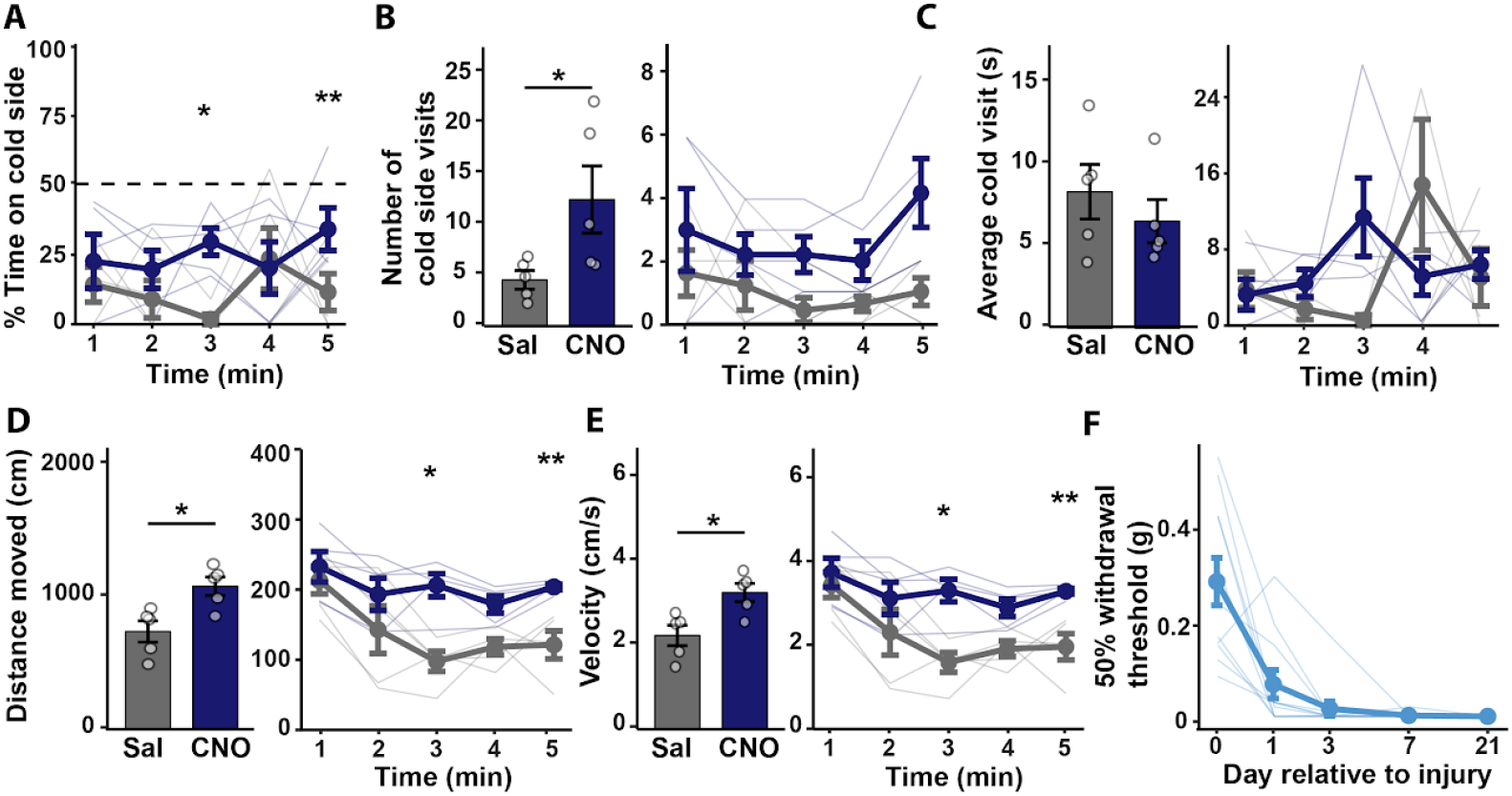
Cold place avoidance extended data for mice with SNI. **A)** Percentage of time the mice spent on the cold side, minute by minute. **B)** Number of cold side entries for the whole test session (*Left*) and number of cold side entries minute by minute (*Right*). **C)** Average duration of cold side visits (total time on cold side divided by number of visits for the whole test session (*Left*) and average cold side visit duration minute by minute (*Right*). **D)** Total distance traveled over the entire experiment (*Left*) and total distance minute by minute (*Right*). **E)** Average velocity over the entire experiment (*Left*) and average velocity minute by minute (*Right*). **F)** Development of mechanical hypersensitivity by Von Frey up-down testing at multiple time points after SNI surgery. *p < 0.05, **p < 0.01. All error bars indicate SEM.

**Supplementary Figure 9.**
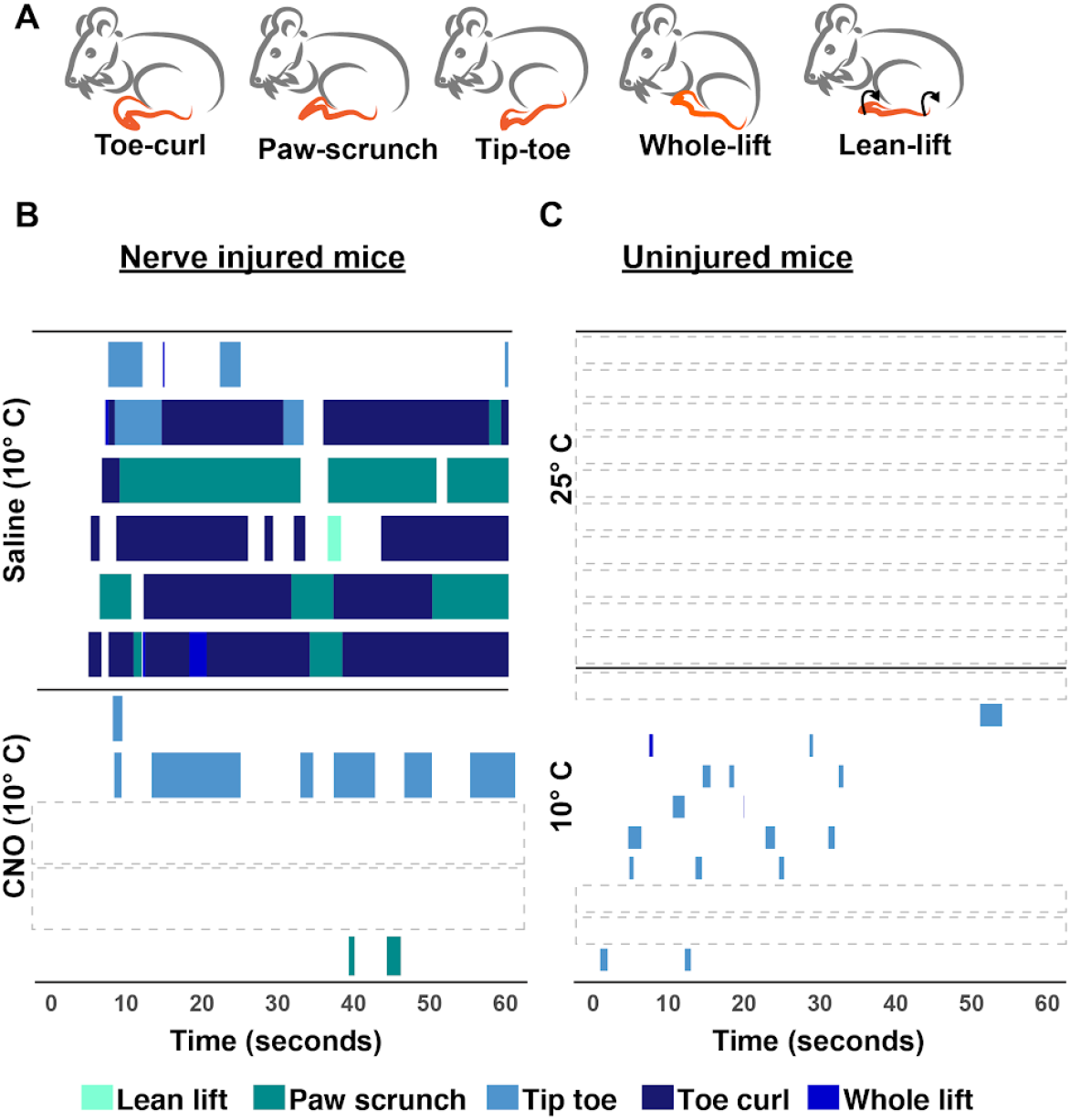
Cold plate extended data. **A)** Visual depiction of the multiple behaviors mice spontaneously show on the cold plate after SNI. **B)** Behavior raster plots of all behaviors all animals displayed on the cold plate after either saline or CNO injection. Each row represents one mouse for a 60-second duration for the test. **C)** Spontaneous behaviors are displayed at either room temperature or cold temperature in uninjured animals.

**Supplementary Figure 10.**
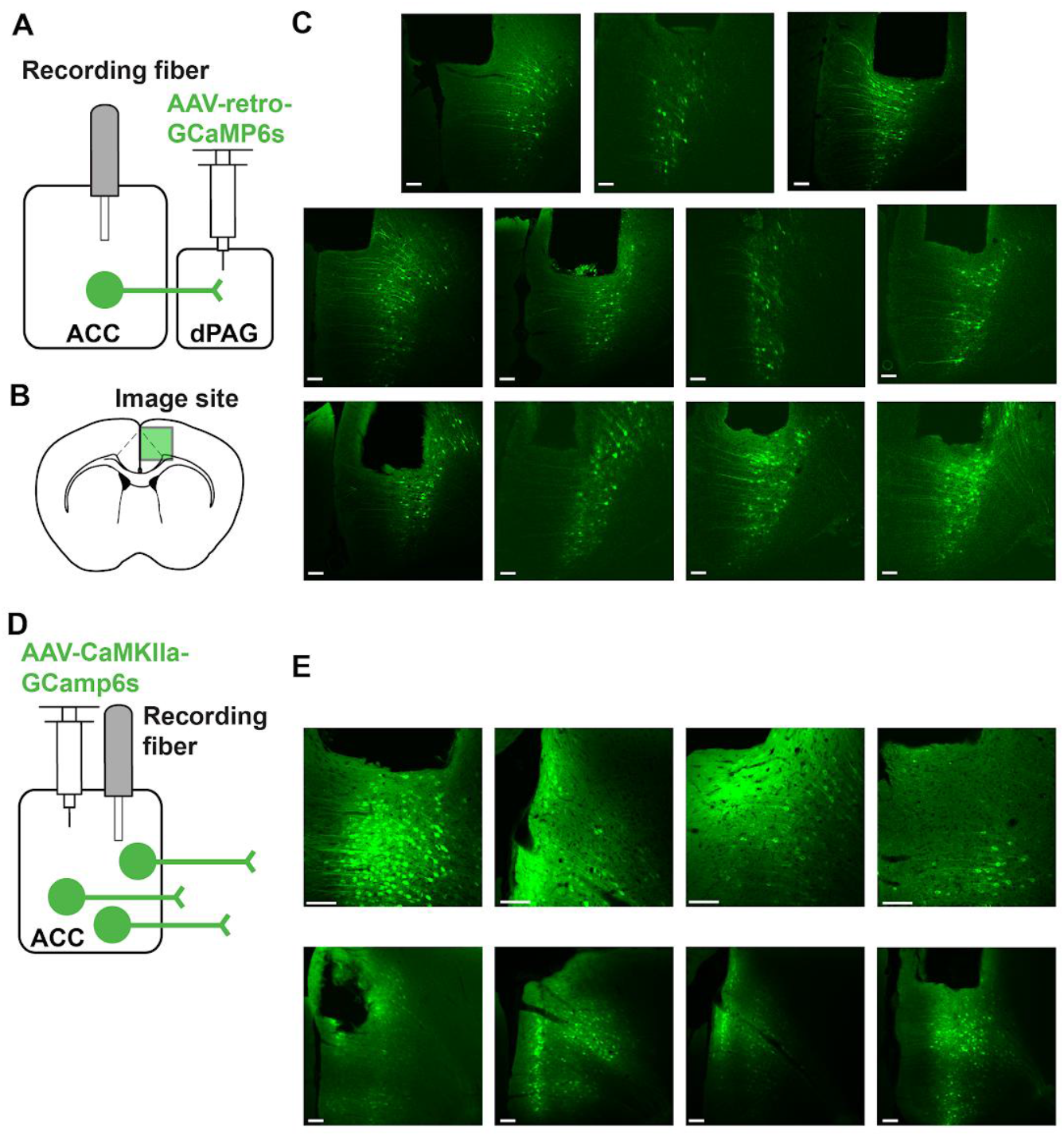
GCaMP6s and fiber placement verification. **A)** Virus injection to record ACC→dlPAG projection neurons. **B)** Schematic depiction of the approximate location at which histological images were taken. **C)** GCaMP6s and fiber damage in the ACC from theACC→dlPAG cohort. **D)** Virus injection to record CaMKIIα ACC pyramidal neurons. **E)** GCaMP6s and fiber damage in the ACC from the CaMKIIα cohort. All scale bars = 100 µm.

**Supplementary Figure 11.**
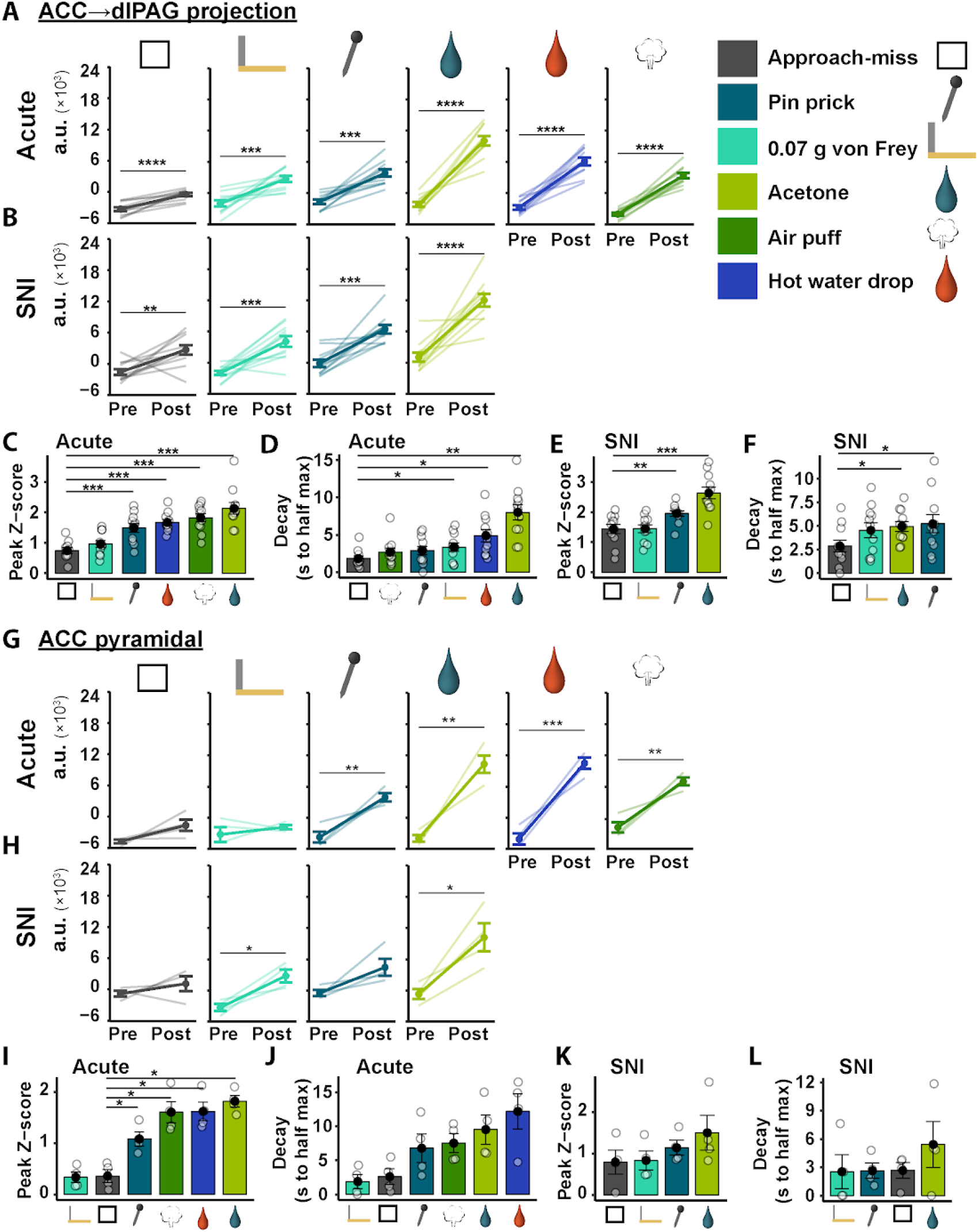
Magnitude and characteristics of photometry responses to punctate stimuli. **A)** Magnitude (area under the curve) of the fluorescence in ACC→dlPAG neurons in the 30-second window pre and post-stimulus. **B)** Magnitude of the fluorescence in ACC→dlPAG neurons in the 30-second window pre and post-stimulus, 3 weeks after SNI. **C)** Peak Z-score in the 30-second post-response window for each stimulus, ranked from lowest (left) to highest (right), in uninjured ACC→dlPAG animals. **D)** Decay measure in the 30-second post-response window for each stimulus, ranked from lowest (*Left*) to highest (*Right*), in uninjured ACC→dlPAG animals. **E)** Peak Z-score in the 30-second post-response window for each stimulus, ranked from lowest (*Left*) to highest (*Right*), 3 weeks after SNI in ACC→dlPAG animals. **F)** Decay measure in the 30-second post-response window for each stimulus, ranked from lowest (*Left*) to highest (*Right*), 3 weeks after SNI in ACC→dlPAG animals. **G)** Magnitude of the fluorescence in CaMKIIα ACC neurons in the 30-second window pre-and post-stimulus, in uninjured animals. **H)** Magnitude of the fluorescence in CaMKIIα ACC neurons in the 30-second window pre and post-stimulus, 3 weeks after SNI. **I)** Peak Z-score in the 30-second post-response window for each stimulus, ranked from lowest (*Left*) to highest (*Right*), in uninjured CaMKIIα ACC animals. **J)** Decay measure in the 30-second post-response window for each stimulus, ranked from lowest (*Left*) to highest (*Right*), in uninjured CaMKIIα animals. **K)** Peak Z-score in the 30-second post-response window for each stimulus, ranked from lowest (*Left*) to highest (*Right*), 3 weeks after SNI in CaMKIIα animals. **L)** Decay measure in the 30-second post-response window for each stimulus, ranked from lowest (*Left*) to highest (*Right*), 3 weeks after SNI in CaMKIIα animals. *p < 0.05, **p < 0.01, ***p < 0.001, ****p < 0.0001. All error bars indicate SEM.

**Supplementary Figure 12.**
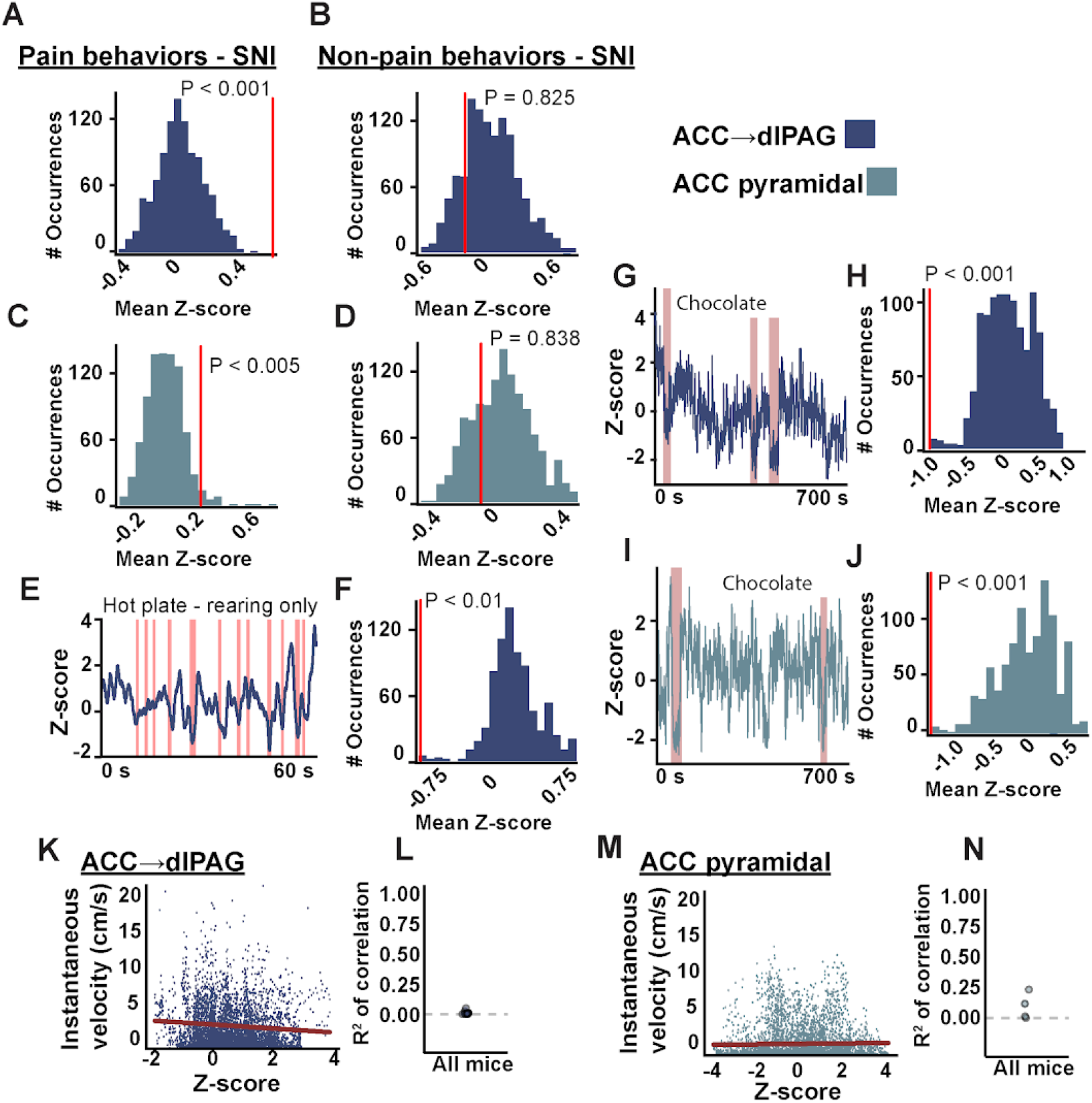
ACC photometry behavior testing extended data. **A)** Representative histogram from a permutation test for pain behaviors in anACC→dlPAG mouse 3 weeks after SNI. **B)** Representative histogram from a permutation test for non-pain behaviors in an ACC→dlPAG mouse 3 weeks after SNI. **C)** Representative histogram from a permutation test for pain behaviors in a CaMKIIα mouse 3 weeks after SNI. **D)** Representative histogram from a permutation test for non-pain behaviors in a CaMKIIα ACC mouse 3 weeks after SNI. **E)** Hot plate representative trace from an ACC→dlPAG recording with transparent red bars overlaying rearing behavioral epochs. **F)** Representative histogram from a permutation test for rearing behavior epochs from recording in E. Not all P-values were below 0.05, but there was a trend of low P-values. **G)** Representative trace from an ACC→dlPAG neuron recording in an uninjured animal free foraging for pieces of chocolate. Overlaying transparent brown bars correspond to epochs of holding and eating a piece of chocolate. **H)** Representative permutation test for chocolate consumption in G. **I)** Representative trace from a CaMKIIα recording in an uninjured animal, free foraging for pieces of chocolate. **J)** Representative permutation test for chocolate consumption in I. **K)** Representative plot of the relationship between ACC→dlPAG fluorescence and instantaneous velocity for one mouse. Each colored dot represents a frame from an open-field recording session. The red line is linear regression. **L)** Pearson correlation scores for all ACC→dlPAG mice. **M)** Representative plot of the relationship between CaMKIIα ACC fluorescence and instantaneous velocity. **N)** Pearson correlation scores for all CaMKIIα mice.

## References

Acuña MA, Kasanetz F, De Luna P, Falkowska M, Nevian T. 2023. Principles of nociceptive coding in the anterior cingulate cortex. Proc Natl Acad Sci U S A 120:e2212394120.

Apkarian AV, Bushnell MC, Treede R-D, Zubieta J-K. 2005. Human brain mechanisms of pain perception and regulation in health and disease. Eur J Pain 9:463–484.

Bagley EE, Ingram SL. 2020. Endogenous opioid peptides in the descending pain modulatory circuit. Neuropharmacology 173:108131.

Baker A, Kalmbach B, Morishima M, Kim J, Juavinett A, Li N, Dembrow N. 2018. Specialized subpopulations of deep-layer pyramidal neurons in the neocortex: Bridging cellular properties to functional consequences. J Neurosci 38:5441–5455.

Ballantine HT Jr, Cassidy WL, Flanagan NB, Marino R Jr. 1967. Stereotaxic anterior cingulotomy for neuropsychiatric illness and intractable pain. J Neurosurg 26:488–495.

Bandler R, Shipley MT. 1994. Columnar organization in the midbrain periaqueductal gray: modules for emotional expression? Trends Neurosci 17:379–389.

Baron R. 2000. Peripheral neuropathic pain: from mechanisms to symptoms. Clin J Pain 16:S12– 20.

Barthas F, Sellmeijer J, Hugel S, Waltisperger E, Barrot M, Yalcin I. 2015. The anterior cingulate cortex is a critical hub for pain-induced depression. Biol Psychiatry 77:236–245.

Basbaum AI, Fields HL. 1978. Endogenous pain control mechanisms: review and hypothesis. Ann Neurol 4:451–462.

Behbehani MM. 1995. Functional characteristics of the midbrain periaqueductal gray. Prog Neurobiol 46:575–605.

BRAIN Initiative Cell Census Network (BICCN). 2021. A multimodal cell census and atlas of the mammalian primary motor cortex. Nature 598:86–102.

Bushnell MC, Čeko M, Low LA. 2013. Cognitive and emotional control of pain and its disruption in chronic pain. Nat Rev Neurosci 14:502–511.

Calejesan AA, Kim SJ, Zhuo M. 2000. Descending facilitatory modulation of a behavioral nociceptive response by stimulation in the adult rat anterior cingulate cortex. Eur J Pain 4:83– 96.

Carter CS, Botvinick MM, Cohen JD. 1999. The contribution of the anterior cingulate cortex to executive processes in cognition. Reviews in the Neurosciences. doi:10.1515/REVNEURO.1999.10.1.49

Chen T, Koga K, Descalzi G, Qiu S, Wang J, Zhang L-S, Zhang Z-J, He X-B, Qin X, Xu F-Q, Hu J, Wei F, Huganir RL, Li Y-Q, Zhuo M. 2014. Postsynaptic potentiation of corticospinal projecting neurons in the anterior cingulate cortex after nerve injury. Mol Pain 10:33.

Chen T, Taniguchi W, Chen QY, Tozaki-Saitoh H, Song Q, Liu RH, Koga K, Matsuda T, Kaito-Sugimura Y, Wang J, Li ZH, Lu YC, Inoue K, Tsuda M, Li YQ, Nakatsuka T, Zhuo M. 2018. Top-down descending facilitation of spinal sensory excitatory transmission from the anterior cingulate cortex. Nat Commun 9. doi:10.1038/s41467-018-04309-2

Choi J, Lee Y-B, So D, Kim JY, Choi S, Kim S, Keum S. 2025. Cortical representations of affective pain shape empathic fear in male mice. Nat Commun 16:1937.

Cohen RA, Paul R, Zawacki TM, Moser DJ, Sweet L, Wilkinson H. 2001. Emotional and personality changes following cingulotomy. Emotion 1:38–50.

Corder G, Ahanonu B, Grewe BF, Wang D, Schnitzer MJ, Scherrer G. 2019. An amygdalar neural ensemble that encodes the unpleasantness of pain. Science 363:276–281.

Corder G, Tawfik VL, Wang D, Sypek EI, Low SA, Dickinson JR, Sotoudeh C, Clark JD, Barres BA, Bohlen CJ, Scherrer G. 2017. Loss of μ opioid receptor signaling in nociceptors, but not microglia, abrogates morphine tolerance without disrupting analgesia. Nat Med. doi:10.1038/nm.4262

Cox JJ, Reimann F, Nicholas AK, Thornton G, Roberts E, Springell K, Karbani G, Jafri H, Mannan J, Raashid Y, Al-Gazali L, Hamamy H, Valente EM, Gorman S, Williams R, McHale DP, Wood JN, Gribble FM, Woods CG. 2006. An SCN9A channelopathy causes congenital inability to experience pain. Nature 444:894–898.

Del Rio D, Beucher B, Lavigne M, Wehbi A, Gonzalez Dopeso-Reyes I, Saggio I, Kremer EJ. 2019. CAV-2 vector development and gene transfer in the central and peripheral nervous systems. Front Mol Neurosci 12:71.

DeNardo LA, Liu CD, Allen WE, Adams EL, Friedmann D, Fu L, Guenthner CJ, Tessier-Lavigne M, Luo L. 2019. Temporal evolution of cortical ensembles promoting remote memory retrieval. Nat Neurosci 22:460–469.

Deng H, Xiao X, Wang Z. 2016. Periaqueductal gray neuronal activities underlie different aspects of defensive behaviors. J Neurosci 36:7580–7588.

De Ridder D, Adhia D, Vanneste S. 2021. The anatomy of pain and suffering in the brain and its clinical implications. Neurosci Biobehav Rev 130:125–146.

Esmaeilou Y, Tamaddonfard E, Erfanparast A, Soltanalinejad-Taghiabad F. 2022. Behavioral and receptor expression studies on the primary somatosensory cortex and anterior cingulate cortex oxytocin involvement in modulation of sensory and affective dimensions of neuropathic pain induced by partial sciatic nerve ligation in rats. Physiol Behav 251:113818.

Evans DA, Stempel AV, Vale R, Ruehle S, Lefler Y, Branco T. 2018. A synaptic threshold mechanism for computing escape decisions. Nature 558:590–594.

Falconi-Sobrinho LL, Anjos-Garcia TD, Rebelo MA, Hernandes PM, Almada RC, Tanus-Santos JE, Coimbra NC. 2024. The anterior cingulate cortex and its interface with the dorsal periaqueductal grey regulating nitric oxide-mediated panic-like behaviour and defensive antinociception. Neuropharmacology 245:109831.

Falkner AL, Wei D, Song A, Watsek LW, Chen I, Chen P, Feng JE, Lin D. 2020. Hierarchical representations of aggression in a hypothalamic-midbrain circuit. Neuron 106:637–648.e6.

Foltz EL, White LE. 1962. Pain “Relief” by Frontal Cingulumotomy. Journal of Neurosurgery. doi:10.3171/jns.1962.19.2.0089

Franciosa F, Acuña MA, Nevian NE, Nevian T. 2024. A cellular mechanism contributing to pain-induced analgesia. Pain 165:2517–2529.

François A, Low SA, Sypek EI, Christensen AJ, Sotoudeh C, Beier KT, Ramakrishnan C, Ritola KD, Sharif-Naeini R, Deisseroth K, Delp SL, Malenka RC, Luo L, Hantman AW, Scherrer G. 2017. A Brainstem-Spinal Cord Inhibitory Circuit for Mechanical Pain Modulation by GABA and Enkephalins. Neuron 93:822–839.e6.

Franklin TB, Silva BA, Perova Z, Marrone L, Masferrer ME, Zhan Y, Kaplan A, Greetham L, Verrechia V, Halman A, Pagella S, Vyssotski AL, Illarionova A, Grinevich V, Branco T, Gross CT. 2017. Prefrontal cortical control of a brainstem social behavior circuit. Nat Neurosci 20:260–270.

Grahek N. 2012. Feeling Pain and Being in Pain, 2nd ed, A Bradford Book. Cambridge, MA: Bradford Books.

Guenthner CJ, Miyamichi K, Yang HH, Heller HC, Luo L. 2013. Permanent genetic access to transiently active neurons via TRAP: Targeted recombination in active populations. Neuron 79:1257.

Gu L, Uhelski ML, Anand S, Romero-Ortega M, Kim Y-T, Fuchs PN, Mohanty SK. 2015. Pain inhibition by optogenetic activation of specific anterior cingulate cortical neurons. PLoS One 10:e0117746.

Heinricher MM, Cheng ZF, Fields HL. 1987. Evidence for two classes of nociceptive modulating neurons in the periaqueductal gray. Journal of Neuroscience 7:271–278.

Huang T, Lin SH, Malewicz NM, Zhang Y, Zhang Y, Goulding M, LaMotte RH, Ma Q. 2019. Identifying the pathways required for coping behaviours associated with sustained pain. Nature 565:86–90.

Hutchison WD, Davis KD, Lozano AM, Tasker RR, Dostrovsky JO. 1999. Pain-related neurons in the human cingulate cortex. Nat Neurosci 2:403–405.

Iwata K, Kamo H, Ogawa A, Tsuboi Y, Noma N, Mitsuhashi Y, Taira M, Koshikawa N, Kitagawa J. 2005. Anterior cingulate cortical neuronal activity during perception of noxious thermal stimuli in monkeys. J Neurophysiol 94:1980–1991.

Johansen JP, Fields HL, Manning BH. 2001. The affective component of pain in rodents: direct evidence for a contribution of the anterior cingulate cortex. Proc Natl Acad Sci U S A 98:8077–8082.

Journée SH, Mathis VP, Fillinger C, Veinante P, Yalcin I. 2023. Janus effect of the anterior cingulate cortex: Pain and emotion. Neuroscience & Biobehavioral Reviews 153:105362. doi:10.1016/j.neubiorev.2023.105362

Keay KA, Bandler R. 2015. Periaqueductal GrayThe Rat Nervous System. Elsevier. pp. 207–221.

Kim EJ, Horovitz O, Pellman BA, Tan LM, Li Q, Richter-Levin G, Kim JJ. 2013. Dorsal periaqueductal gray-amygdala pathway conveys both innate and learned fear responses in rats. Proc Natl Acad Sci U S A 110:14795–14800.

Koga K, Descalzi G, Chen T, Ko HG, Lu J, Li S, Son J, Kim TH, Kwak C, Huganir RL, Zhao MG, Kaang BK, Collingridge GL, Zhuo M. 2015. Coexistence of two forms of LTP in ACC provides a synaptic mechanism for the interactions between anxiety and chronic pain. Neuron 85:377– 389.

Kuner R, Tan LL. 2021. Neocortical circuits in pain and nociception. Nature Reviews Neuroscience 22:458–471. doi:10.1038/s41583-021-00468-2.

Kuo C-C, Yen C-T. 2005. Comparison of anterior cingulate and primary somatosensory neuronal responses to noxious laser-heat stimuli in conscious, behaving rats. J Neurophysiol 94:1825– 1836.

LaGraize SC, Labuda CJ, Rutledge MA, Jackson RL, Fuchs PN. 2004. Differential effect of anterior cingulate cortex lesion on mechanical hypersensitivity and escape/avoidance behavior in an animal model of neuropathic pain. Exp Neurol 188:139–148.

Lee J-Y, You T, Lee C-H, Im GH, Seo H, Woo C-W, Kim S-G. 2022. Role of anterior cingulate cortex inputs to periaqueductal gray for pain avoidance. Curr Biol 32:2834–2847.e5.

Livrizzi G, Chang-Weinberg J, Johnson DA, Lubejko ST, Liao J, Kimmey BA, Dong C, Li Y, Beier KT, Corder G, Tian L, Banghart MR. 2026. Top-down control of the descending pain modulatory system drives multimodal placebo analgesia. Neuron. doi:10.1016/j.neuron.2026.03.025

Li X-Y, Ko H-G, Chen T, Descalzi G, Koga K, Wang H, Kim SS, Shang Y, Kwak C, Park S-W, Shim J, Lee K, Collingridge GL, Kaang B-K, Zhuo M. 2010. Alleviating neuropathic pain hypersensitivity by inhibiting PKMzeta in the anterior cingulate cortex. Science 330:1400– 1404.

López-González MV, González-García M, Peinado-Aragonés CA, Barbancho MÁ, Díaz-Casares A, Dawid-Milner MS. 2020. Pontine A5 region modulation of the cardiorespiratory response evoked from the midbrain dorsolateral periaqueductal grey. J Physiol Biochem 76:561–572.

Manglik A, Lin H, Aryal DK, McCorvy JD, Dengler D, Corder G, Levit A, Kling RC, Bernat V, Hübner H, Huang XP, Sassano MF, Giguère PM, Löber S, Duan D, Scherrer G, Kobilka BK, Gmeiner P, Roth BL, Shoichet BK. 2016. Structure-based discovery of opioid analgesics with reduced side effects. Nature. doi:10.1038/nature19112

Ma X, Yu W, Yao P’an, Zhu Y, Dai J, He X, Liu B, Xu C, Shao X, Fang J, Shen Z. 2022. Afferent and efferent projections of the rostral anterior cingulate cortex in young and middle-aged mice. Front Aging Neurosci 14:960868.

Oswell CS, Rogers SA, James JG, McCall NM, Hsu AI, Salimando GJ, Mahmood M, Wooldridge LM, Wachira M, Jo AY, Sandoval Ortega RA, Wojick JA, Beattie K, Farinas SA, Chehimi SN, Rodrigues A, Wu JWK, Ejoh LL, Kimmey BA, Lo E, Azouz G, Vasquez JJ, Banghart MR, Beier KT, Creasy KT, Crist RC, Ramakrishnan C, Reiner BC, Deisseroth K, Yttri EA, Corder G. 2026. Mimicking opioid analgesia in cortical pain circuits. Nature 649:938–947.

Qu C, King T, Okun A, Lai J, Fields HL, Porreca F. 2011. Lesion of the rostral anterior cingulate cortex eliminates the aversiveness of spontaneous neuropathic pain following partial or complete axotomy. Pain 152:1641–1648.

Rainville P, Duncan GH, Price DD, Carrier B, Bushnell MC. 1997. Pain affect encoded in human anterior cingulate but not somatosensory cortex. Science 277:968–971.

Reis FMCV, Liu J, Schuette PJ, Lee JY, Maesta-Pereira S, Chakerian M, Wang W, Canteras NS, Kao JC, Adhikari A. 2021. Shared dorsal periaqueductal gray activation patterns during exposure to innate and conditioned threats. J Neurosci 41:5399–5420.

Reis FM, Lee JY, Maesta-Pereira S, Schuette PJ, Chakerian M, Liu J, La-Vu MQ, Tobias BC, Ikebara JM, Kihara AH, Canteras NS, Kao JC, Adhikari A. 2021. Dorsal periaqueductal gray ensembles represent approach and avoidance states. Elife 10. doi:10.7554/eLife.64934

Roth BL. 2016. DREADDs for Neuroscientists. Neuron. doi:10.1016/j.neuron.2016.01.040

Santello M, Nevian T. 2015. Dysfunction of cortical dendritic integration in neuropathic pain reversed by serotoninergic neuromodulation. Neuron 86:233–246.

Shi W, Xue M, Wu F, Fan K, Chen Q-Y, Xu F, Li X-H, Bi G-Q, Lu J-S, Zhuo M. 2022. Whole-brain mapping of efferent projections of the anterior cingulate cortex in adult male mice. Mol Pain 18:17448069221094529.

Shyu B-C, Chen W-F, Shih H-C. 2008. Electrically and mechanically evoked nociceptive neuronal responses in the rat anterior cingulate cortex. Acta Neurochir Suppl 101:23–25.

Sikes RW, Vogt BA. 1992. Nociceptive neurons in area 24 of rabbit cingulate cortex. J Neurophysiol 68:1720–1732.

Smith ML, Asada N, Malenka RC. 2021. Anterior cingulate inputs to nucleus accumbens control the social transfer of pain and analgesia. Science 371:153–159.

Tasic B, Yao Z, Graybuck LT, Smith KA, Nguyen TN, Bertagnolli D, Goldy J, Garren E, Economo MN, Viswanathan S, Penn O, Bakken T, Menon V, Miller J, Fong O, Hirokawa KE, Lathia K, Rimorin C, Tieu M, Larsen R, Casper T, Barkan E, Kroll M, Parry S, Shapovalova NV, Hirschstein D, Pendergraft J, Sullivan HA, Kim TK, Szafer A, Dee N, Groblewski P, Wickersham I, Cetin A, Harris JA, Levi BP, Sunkin SM, Madisen L, Daigle TL, Looger L, Bernard A, Phillips J, Lein E, Hawrylycz M, Svoboda K, Jones AR, Koch C, Zeng H. 2018. Shared and distinct transcriptomic cell types across neocortical areas. Nature. doi:10.1038/s41586-018-0654-5

Tervo DGR, Hwang BY, Viswanathan S, Gaj T, Lavzin M, Ritola KD, Lindo S, Michael S, Kuleshova E, Ojala D, Huang CC, Gerfen CR, Schiller J, Dudman JT, Hantman AW, Looger LL, Schaffer DV, Karpova AY. 2016. A Designer AAV Variant Permits Efficient Retrograde Access to Projection Neurons. Neuron. doi:10.1016/j.neuron.2016.09.021

Tjølsen A, Berge O-G, Hunskaar S, Rosland JH, Hole K. 1992. The formalin test: an evaluation of the method. Pain 51:5–17.

Treede RD, Kenshalo DR, Gracely RH, Jones AKP. 1999. The cortical representation of pain. Pain 79:105–111.

Vander Weele CM, Siciliano CA, Matthews GA, Namburi P, Izadmehr EM, Espinel IC, Nieh EH, Schut EHS, Padilla-Coreano N, Burgos-Robles A, Chang C-J, Kimchi EY, Beyeler A, Wichmann R, Wildes CP, Tye KM. 2018. Dopamine enhances signal-to-noise ratio in cortical-brainstem encoding of aversive stimuli. Nature 563:397–401.

Vaughn E, Eichhorn S, Jung W, Zhuang X, Dulac C. 2022. Three-dimensional interrogation of cell types and instinctive behavior in the periaqueductal gray. bioRxiv. doi:10.1101/2022.06.27.497769

Vogt BA. 2005. Pain and emotion interactions in subregions of the cingulate gyrus. Nat Rev Neurosci 6:533–544.

Wager TD, Atlas LY, Lindquist MA, Roy M, Woo C-W, Kross E. 2013. An fMRI-based neurologic signature of physical pain. N Engl J Med 368:1388–1397.

Wang H, Flores RJ, Yarur HE, Limoges A, Bravo-Rivera H, Casello SM, Loomba N, Enriquez-Traba J, Arenivar M, Wang Q, Ganley R, Ramakrishnan C, Fenno LE, Kim Y, Deisseroth K, Or G, Dong C, Hoon MA, Tian L, Tejeda HA. 2024. Prefrontal cortical dynorphin peptidergic transmission constrains threat-driven behavioral and network states. Neuron 112:2062– 2078. doi:10.1016/j.neuron.2024.03.015

Wang J-Y, Luo F, Chang J-Y, Woodward DJ, Han J-S. 2003. Parallel pain processing in freely moving rats revealed by distributed neuron recording. Brain Res 992:263–271.

Wang J, Zhang X, Cao B, Liu J, Li Y. 2015. Facilitation of synaptic transmission in the anterior cingulate cortex in viscerally hypersensitive rats. Cereb Cortex 25:859–868.

Wei F, Zhuo M. 2001. Potentiation of sensory responses in the anterior cingulate cortex following digit amputation in the anaesthetised rat. J Physiol 532:823–833.

